# Epistasis Arises from Shifting the Rate-Limiting Step during Enzyme Evolution

**DOI:** 10.1101/2023.06.29.547057

**Authors:** Christopher Fröhlich, H. Adrian Bunzel, Karol Buda, Adrian J. Mulholland, Marc W. van der Kamp, Pål J. Johnsen, Hanna-Kirsti S. Leiros, Nobuhiko Tokuriki

## Abstract

The molecular mechanisms by which epistasis boosts enzyme activity remain elusive, undermining our ability to predict the evolution of pathogens and engineer novel biocatalysts. Here, we reveal how directed evolution of a β-lactamase yielded highly epistatic activity enhancements. Evolution selected four mutations that increase antibiotic resistance 40-fold, despite their marginal individual effects (≤ 2-fold). Synergistic improvements coincided with the introduction of super-stochiometric burst kinetics, indicating that epistasis is rooted in the enzyme’s conformational dynamics. Kinetic, structural, and dynamical analyses reveal that epistasis was driven by distinct effects of each mutation on the catalytic cycle. The first mutation acquired during evolution increases protein flexibility and accelerates substrate binding, which is rate-limiting in the wild-type enzyme. The ensuing mutations predominantly boosted the chemical steps by fine-tuning substrate interactions. Our work identifies an overlooked cause for epistasis: changing the rate-limiting step can result in substantial positive synergy boosting enzyme activity.

## INTRODUCTION

Enzymes rely on intricate intramolecular interactions between amino acids to organize the active site and achieve efficient catalysis. Rewiring these interactions through evolution often leads to unexpected and non-additive effects on protein fitness in a phenomenon known as epistasis.^1–4^ Positive epistasis, wherein the effect of mutations is more beneficial when combined than in isolation, is highly desirable in protein engineering and often drives evolutionary trajectories. In contrast, negative epistasis is caused by mutations that act antagonistically, which adversely affects protein fitness and can be detrimental to protein engineering campaigns. Consequently, epistasis often dictates enzyme evolution by providing access to, or restricting, mutational paths.^1–6^ Such non-additive interactions can be introduced by various factors, including changes in structural interactions and protein conformational dynamics. For example, epistasis can arise by rewiring the interactions of active site residues, thereby establishing new interactions with the substrate. Also, mutations that change conformational dynamics can improve enzymatic activity in a highly synergistic fashion,^7–11^ for instance, by epistatically altering the dynamics of solvent-exposed loops to aid substrate entry or enhance active-site organization.^12–14^

While the link between structural changes and epistasis has been intensively studied, the mechanistic relationship between epistatic mutations and the overall catalytic cycle is often overlooked. Concerning epistasis, mechanistic studies may be particularly important because evolution typically enhances the slowest steps in the catalytic cycle,^15–18^ which might change the rate-limiting step resulting in non-additive effects. Here, we hypothesize that studying how mutations impact each step in the reaction could reveal novel mechanisms of epistasis, which would improve the predictability of evolution and provide a better understanding of the overall permissiveness of adaptive landscapes.

The β-lactamase OXA-48 is an excellent model system for studying epistasis in the context of antimicrobial resistance development.^19–21^ OXA-48 uses a catalytic serine (S70) to cleave β-lactams in a three-step mechanism comprising enzyme-substrate complex formation (ES), formation of an acyl-enzyme intermediate (EI), and hydrolytic product release (E+P).^22^ OXA-48 confers resistance to many carbapenem and penicillin β-lactams, but only slowly hydrolyses oxyimino-cephalosporins such as ceftazidime (CAZ).^23^ Low catalytic activity for CAZ hydrolysis has been attributed to the substrate’s bulkiness, potentially requiring sampling of alternative loop conformations to promote cephalosporin binding and hydrolysis.^12, 24^ While we have recently demonstrated that single mutations in OXA-48 can result in low-level CAZ resistance by increasing the flexibility of active site loops,^19–21^ the mutational effects on the overall catalytic cycle and their potential for epistasis remain elusive.

Here, we used directed evolution to study the mechanistic drivers of epistasis during the adaptation of an antimicrobial resistance gene. After subjecting the β-lactamase OXA-48 to iterative rounds of mutagenesis and selection, we constructed an adaptive fitness landscape of the introduced mutations that identifies positive epistasis as a key driver for enzyme evolution. By combining biochemical, structural, and computational methods, we reveal how evolution alters the protein’s conformational dynamics to accelerate substrate binding and hydrolysis, thereby changing the rate-limiting step and introducing epistasis. This detailed understanding of mutational effects on the whole catalytic cycle is crucial to predicting epistatic interactions^1–4^, which is relevant to the evolution of enzymatic activity and the design and engineering of novel enzymes.^25^

## RESULTS

### Evolution of OXA-48 is driven by positive epistasis

To investigate how epistasis drives the evolution of OXA-48, we performed five cycles of directed evolution starting from the wild-type OXA-48 (wtOXA-48), using error-prone PCR mutagenesis followed by selection on increasing CAZ concentrations (Fig. 1a). Variants arising along the evolutionary trajectory were characterized by their half-maximal inhibitory concentrations (*IC_50_*) from antibiotic dose-response growth curves (Tab. S1). Five mutations were accumulated during evolution (F72L→S212A→T213A→A33V→K51E), resulting in Q5 that confers 43-fold increased CAZ resistance in *E. coli* (Fig. 1b, Tab. S1). The mutations acquired during evolution cluster around the active site or structural elements known to alter substrate specificity, such as the nucleophilic S70 or the Ω- and β5-β6 loops (Fig. 1c).^23^ K51E increased resistance development by only 1.1-fold compared to Q4. Thus, we focused analysis on the molecular origins of epistasis in Q4 (A33V/F72L/S212A/T213A), which conferred a 40-fold higher CAZ resistance over wtOXA-48 in *E. coli* (Tab. S1).

To study the interplay between mutations acquired along the evolutionary trajectory, we constructed an adaptive landscape of all 16 mutational combinations based on the four mutations in Q4 and determined their *IC*_50_ values when produced in *E. coli* (Fig. 1d, Tab. S2). Overall, positive epistasis greatly shaped the evolution of Q4 and resulted in a resistance increase of 40-fold in contrast to the 3.4-fold predicted increase for strictly additive gains from single point mutations (Fig. 1e, Fig. S1a). To quantify the apparent positive epistasis, we analyzed the contribution of each mutation to the *IC*_50_ fold-change across every possible genetic background (Fig. S1b). Strikingly, epistasis is primarily driven by interactions with F72L, the first mutation acquired during evolution and the only single-point mutation that significantly (2-fold) increased resistance in the wild-type background (analysis of variance [ANOVA], df=4, *P* < 0.001, Tab. S3). For example, the combination of F72L with either S212A or T213A (referred to as alanine mutations hereafter) confers 8.2-fold and 11.7-fold higher resistance than expected (Fig. S1c). We note that changes in thermostability (*T*_m_) did not drive epistatic adaptation (Fig. S1d, Tab. S4). Except for F72L, which reduced the *T*_m_ by 6°C, the selected mutations barely affected the thermostability. Thus, the observed epistasis does not stem from changes in protein stability, but from specific intramolecular interactions that affect the enzyme’s function. Interestingly, we observed diminishing return epistasis in resistance development for higher-order variants along the evolutionary trajectory (Fig. 1f). As described above, F72L increased the *IC*_50_ by 2-fold, followed by a 5-fold gain in *IC*_50_ through S212A. All later mutations, however, provided lower overall improvements. For instance, addition of T213A to F72L/S212A, resulting in Q3, showed diminishing returns and only conferred a 3- fold increase in resistance, which is lower than expected based on the 6.3-fold effect of T213A in F72L. Given that the alanine mutations are located in neighboring positions, they probably have redundant effects on the CAZ *IC*_50_. Thus, their effects do not combine additively and lead to lower-than-expected improvements.

**Figure 1:**
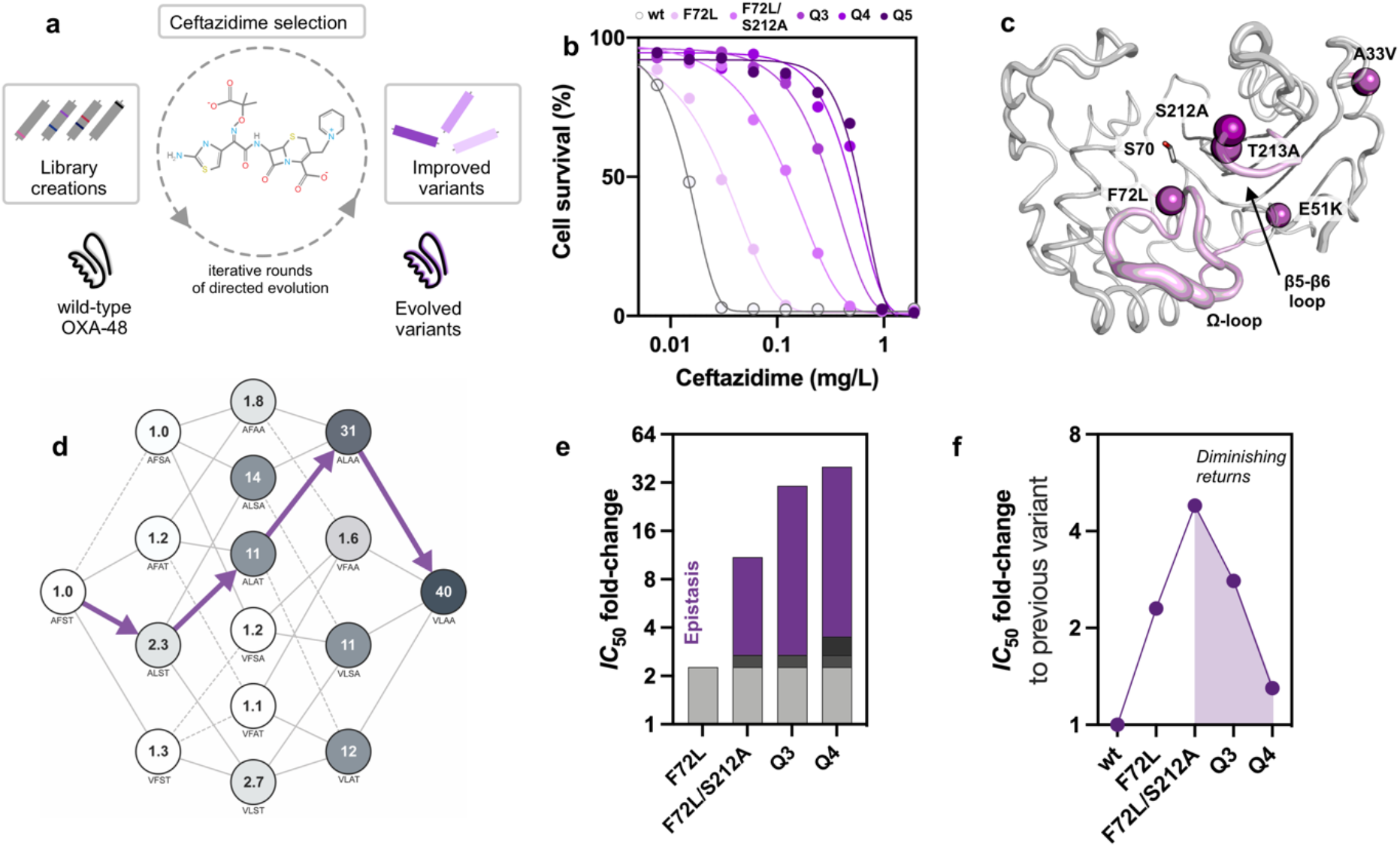
Positive epistasis drives the evolution of OXA-48. **a.** During directed evolution of OXA-48, selection for resistance against the oxyimino-cephalosporin ceftazidime (CAZ) was performed at increasing CAZ concentrations from 0.5 to 14 µM. **b.** CAZ resistance conferred by OXA-48 was improved 43-fold over five rounds of evolution. **c.** Mutations acquired during evolution, such as F72L, S212, and T213A, cluster around the active site serine (S70) and the Ω- and β5-β6 loops that affect substrate specificity (purple). **d.** The adaptive landscape of the mutations found during evolution shows high epistasis. Each node represents a unique variant indicated by single-letter amino acid codes. Values within each node reflect the CAZ *IC50* fold-change relative to wtOXA-48. Purple arrows indicate the trajectory followed during evolution. **e.** Comparison of the effects of single mutations (grey = F72L, dark grey = S212A, black = A33V, no expected effect of T213A) on the *IC50* fold-changes along the evolutionary trajectory reveals a high degree of epistasis (purple). **f.** Comparison of the fold-change improvements relative to the previous variants reveals diminishing returns (purple area) in CAZ resistance.

### Evolution selects for an epistatic burst phase

To understand the molecular origins of epistasis, we studied the reaction kinetics of the OXA- 48 variants (Fig. 2a). We monitored the conversion of CAZ by wtOXA-48, F72L, and Q4 using a stopped-flow and discovered that all enzymes catalyzed the conversion of CAZ with an initial activity burst (Fig. 2b). At 400 µM CAZ, wtOXA-48 possessed a 1.5-fold higher rate in the burst phase than in its subsequent steady-state phase. Interestingly, the burst phase became more pronounced during evolution. F72L and Q4 show 4.6-fold and 48-fold higher burst-phase rates than their respective steady states. Evolution thus selectively improved the burst-phase over the steady-state activity.

To obtain kinetic parameters for the burst phase for a range of variants along the evolutionary trajectory, we assayed substrate conversion with a microtiter plate reader at 4°C (Tab. 1, Fig. S2). As expected, F72L improved the *k*_cat_/*K*_M_ of the burst phase by 8-fold compared to wtOXA-48, while S212A and T213A provided only marginal improvements (∼1.5- fold). Similar to the *IC*_50_ effects, F72L displayed strong pairwise epistasis with either S212A or T213A at the kinetic level and improved *k*_cat_/*K*_M_ by 70-fold and 60-fold compared to wtOXA-48, respectively. Positive epistasis shaped the evolution of *k*_cat_/*K*_M_, culminating in a 470-fold and 800-fold improvement in Q3 and Q4, respectively. The burst-phase *k*_cat_/*K*_M_ values thus show excellent correlation with the *in vivo IC_50_* values (*R*^2^ = 0.97; Pearson correlation, df = 7, *P* < 0.001; Fig. 2c), much better than the correlation between *IC_5_*_0_ and steady-state turnover (*R*^2^ = 0.85; Pearson correlation, df = 7, *P* < 0.001; Fig. S2 to S4, Tab. 1 and S5). Consequently, our kinetic data suggest that evolution selected for burst phases with high epistasis and catalytic efficiency.

**Table 1:** *IC*_50_ and burst-phase catalytic parameters of OXA-48 variants^a,b^.

| | $IC_{50}$<br>(mg/L) | $k_{cat}$<br>(s<sup>-1</sup>) | $K_M$<br>(μM) | $k_{cat}/K_M$<br>(M<sup>-1</sup>s<sup>-1</sup>) |
| --- | --- | --- | --- | --- |
| wtOXA-48 | 0.013 ± 0.002 | - | - | 3.6 |
| F72L | 0.029 ± 0.003 | 0.005 ± 0.001 | 165 ± 15 | 30 |
| S212A | 0.015 ± 0.001 | - | - | 6.0 |
| T213A | 0.013 ± 0.001 | - | - | 5.4 |
| F72L/S212A | 0.140 ± 0.002 | 0.025 ± 0.002 | 95 ± 20 | 260 |
| F72L/T213A | 0.177 ± 0.037 | 0.036 ± 0.009 | 160 ± 70 | 225 |
| S212A/T213A | 0.023 ± 0.003 | - | - | 35 |
| Q3 | 0.389 ± 0.015 | 0.018 ± 0.002 | 10 ± 3 | 1700 |
| Q4 | 0.513 ± 0.054 | 0.077 ± 0.006 | 27 ± 5 | 2900 |

<sup>a</sup> Errors are reported as the standard deviation of the mean.<sup>b</sup> Catalytic parameters were determined at 4°C.
- Not calculated due to linearity of the Michaelis-Menten plot.

The kinetics of the burst phase unveiled fundamentally different catalytic effects between F72L and either the S212A or T213A variants (Fig. 2d and S2). Michaelis-Menten kinetics of wtOXA-48 did not show saturation due to its high *K*_M_ (≫300 µM). Introduction of F72L led to saturation kinetics and substantially decreased the *K*_M_ to 165 ± 15 µM. In contrast, the Michaelis-Menten plots remained linear upon introducing S212A or T213A and only showed marginal increases in *k*_cat_/*K*_M_ (1.6-fold). Interestingly, the double mutants F72L/S212A and F72L/T213A maintained lower *K*_M_ values than F72L while increasing *k*_cat_ by 5 to 7-fold (Tab. 1, Fig. S2). Thus, epistasis between F72L and S212A or T213A probably originates from an interplay between improved binding and catalysis.

Burst phases have been previously observed in β-lactamases, where the fast formation of an acyl-enzyme intermediate during the first turnover is followed by rate-limiting deacylation.^26, 27^ Surprisingly, the burst phases observed here have amplitudes much larger than a single turnover. For example, Q4 displayed a burst-phase amplitude corresponding to 3.8 turnovers at 400 µM CAZ (Fig. 2b). Such super-stoichiometric bursts cannot be explained by a simple two-step reaction mechanism comprising a fast followed by a slow step.^26, 27^ Instead, super-stoichiometric bursts are likely caused by the inactivation of the enzyme during substrate conversion over several catalytic cycles.^26, 27^ Notably, substrate-induced inactivation appeared fully reversible, as indicated by activity assays after incubating Q4 with CAZ (Fig. S5a). In addition, inactivation is not caused by a change in the oligomeric state, as shown by both dynamic light scattering and size exclusion chromatography (Fig. S5b). Thus, turnover apparently triggers a reversible conformational change to a less active state, resulting in the observed burst.

The dependence of the burst phase on the CAZ concentration provides important insights into its role in evolutionary adaptation. With decreasing CAZ concentrations, the rates of the burst and steady-state phases become similar, and the burst phase amplitude decreases. The burst thus became less pronounced at substrate concentrations down to 50 µM at 25°C (Fig. 2b). This is probably because at lower concentrations, substrate-induced inactivation is rarer, and any enzyme that is deactivated has more time to recover to the active state before encountering another substrate molecule. Notably, selection was performed under even lower substrate concentrations (≤14 µM) compared to the *in vitro* kinetic analysis. Thus, the enzymes probably remained predominantly in the burst state under the selection conditions. Taken together, while evolution boosted resistance at the evolutionarily relevant CAZ concentrations, a catalytic bottleneck emerged at high and physiologically irrelevant concentrations that limits activity after the initial burst.

**Figure 2:**
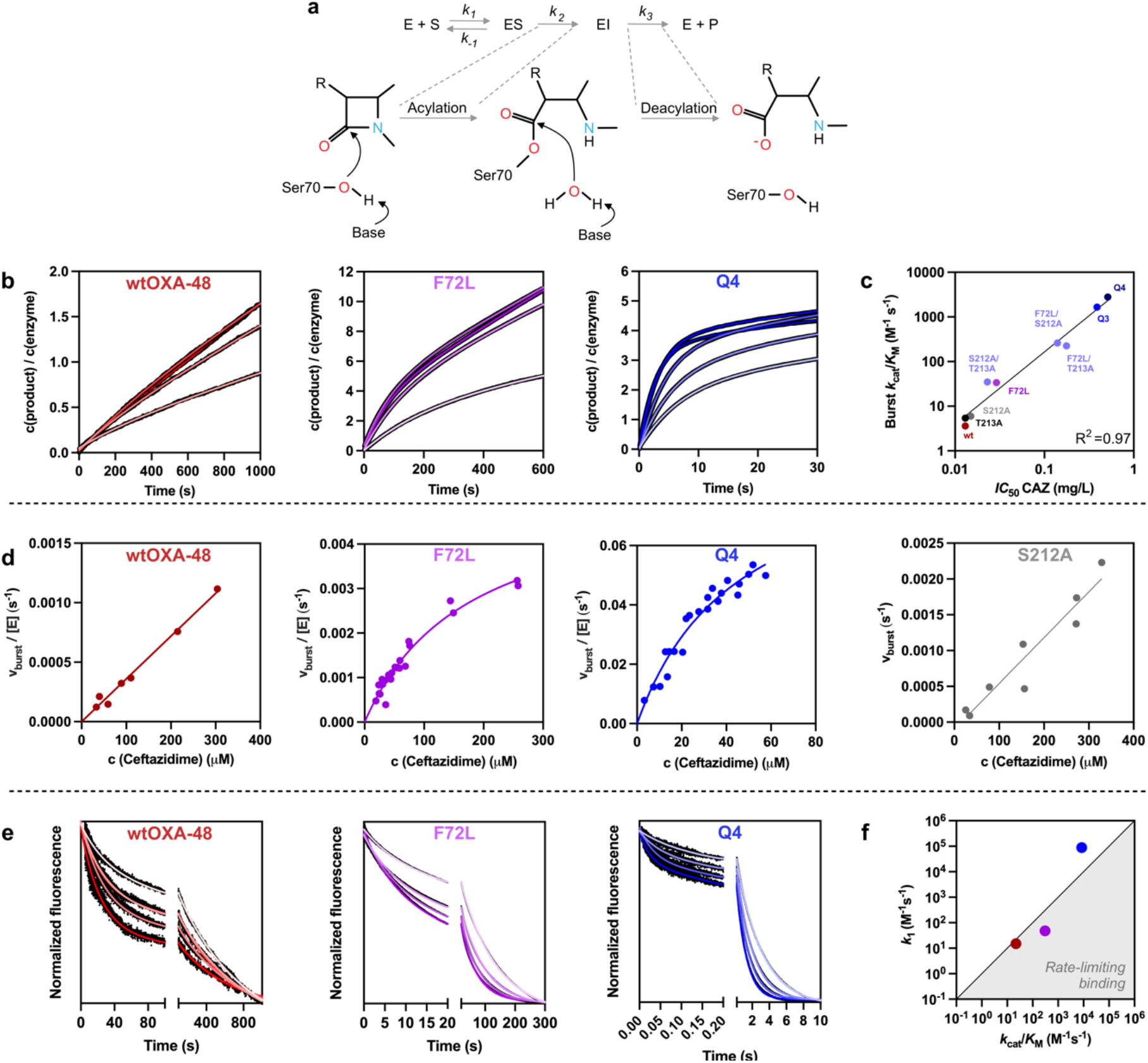
Kinetic changes drive the evolution of OXA-48. **a.** Enzymatic hydrolysis of β-lactams proceeds *via* an enzyme-substrate complex (ES), formation of an acyl-enzyme intermediate (EI), and hydrolytic deacylation (E+P). **b.** Evolution amplified the super-stoichiometric burst behavior of OXA-48 (CAZ concentration: 50 to 400 µM, light to dark colors). **c.** *In vitro* burst phase activities correlate well with the *in vivo IC50* fold changes. **d.** Michaelis-Menten kinetics of the burst phase determined at 4°C**. e.** Substrate binding was measured by W-fluorescence and was substantially accelerated during evolution (CAZ concentration: 100 to 1200 μM, light to dark colors). **f.** Comparison of *k*1 and *k*cat/*K*M between wtOXA-48 (red), F72L (purple), and Q4 (blue) reveals that binding is no longer rate-limiting in the burst phase of Q4 (determined at 25°C, point above the diagonal line at *k*1 =*k*cat/*K*M indicates that binding is not rate-limiting).

### Faster binding and reaction drive positive epistasis

Based on our Michaelis-Menten kinetics, we hypothesized that epistasis in OXA-48 resulted from an interplay between binding and the chemical reaction. F72L apparently unlocked the accessibility of the fitness landscape by substantially accelerating substrate binding, which allowed S212A and T213A to take effect and further boost catalysis (Fig. 2). We dissected this relationship by assaying CAZ binding (*k*_1_ and *k*_-1_, Fig. 2a) and solvent isotope effects in wtOXA-48, F72L, and Q4. By focusing on wtOXA-48 and F72L, we aimed to reveal the mechanistic role of F72L and its ability to recruit subsequent mutations. Comparison to the evolved variant Q4 then allowed us to shed light on the combinatorial effect of F72L with the other mutations.

CAZ binding was assayed by monitoring changes in protein tryptophan fluorescence. In agreement with our burst phase results, CAZ binding progressed with biphasic kinetics (Fig. 2e), where the fast phase probably reflects binding to the burst phase ensemble. The slow phase indicates the presence of a second state that is likely related to the steady-state ensemble. Since the slow phase occurs on a similar timescale as enzyme deactivation and the chemical reaction (Fig. 2b and 2e), it probably reflects a combination of these processes and binding to the less active state. Given that the burst phase is most likely the physiologically relevant phase, we decided to focus our analysis on the fast-binding phase (Tab. 2). For wtOXA-48, CAZ binding (*k*_1_) is 1.5-fold slower than the burst-phase *k*_cat_/*K*_M_ which suggests that binding – and not bond-breaking or product release – limits catalytic efficiency in the wild-type. Notably, *k*_1_ constantly increases during evolution (3.2-fold in F72L and 6000-fold in Q4). In Q4, *k*_1_ is 10-fold faster than *k*_cat_/*K*_M_, indicating that evolution shifted the catalytic bottleneck from substrate binding to either the chemical steps or product release (Fig. 2f). Counterintuitively, the *K*_D_ determined from *k*_-1_/*k*_1_ did not follow the trends observed for *K*_M_ and was 2.6-fold higher for F72L than for wtOXA-48. This observation serves as a reminder that *K*_M_ is an imperfect approximation for substrate affinity and supports our hypothesis that epistasis in OXA-48 has kinetic and not thermodynamic origins. Epistasis is most likely driven by a change in the rate-limiting step resulting from faster substrate binding (*k*_1_) and not by improved substrate affinity (*K*_D_).

**Table 2:** Burst-phase binding and catalysis kinetics.^a^.

|  | wtOXA-48 | F72L | Q4 |
| --- | --- | --- | --- |
| $k_1$ (M<sup>-1</sup> s<sup>-1</sup>) | 15.2 ± 0.5 | 29 ± 3 | 90600 ± 700 |
| $k_{-1}$ (s<sup>-1</sup>) | 0.026 ± 0.001 | 0.127 ± 0.001 | 2.0 ± 0.1 |
| $K_D$ (μM) | 1700 | 4500 | 22 |
| $k_{cat}/K_m$ (M<sup>-1</sup> s<sup>-1</sup>) | 24 | 320 | 9600 |
| Burst-phase KIE<sup>b</sup> | 0.7 | 1.4 | 1.0 |
| Steady-state KIE<sup>b</sup> | 0.8 | 2.8 | 2.5 |

<sup>a</sup> Obtained by stopped-flow at 25°C. Errors are reported as the standard deviation of the mean.<sup>b</sup> determined at 400 μM CAZ.

To further test our hypothesis that evolution shifted the rate-limiting step, we determined solvent isotope effects by assaying product formation at 400 μM CAZ in 80% D_2_O (Tab. 2). Solvent isotope effects >1 indicate rate-limiting deprotonation during the hydrolysis of the acyl-enzyme complex (*k*_3_). The burst-phase isotope effects for wtOXA-48 and Q4 are close to 1, whereas a small isotope effect was observed for F72L (1.4). These isotope effects suggest that hydrolysis of the acyl-enzyme affects the overall rate in F72L, while deacylation is not rate-limiting in wtOXA-48 and Q4. In contrast to the burst-phase effect, F72L and Q4 had pronounced isotope effects of 2.8 and 2.5 in their steady state. Since our data on the evolution of super-stoichiometric burst phases suggest that a conformational change triggers inactivation, the observed increase in isotope effects from the burst phase to the steady state likely indicates that hydrolysis becomes rate-limiting in the less active state. Alternatively, the conformational equilibrium itself could have an isotope effect causing the differences in activity.

Our combined data on binding and activity demonstrate how evolution successively optimized the catalytic cycle and gradually changed the rate-limiting step. In the burst phase, which is probably the physiologically relevant phase, the slowest step appears to be substrate binding in wtOXA-48, binding or deacylation in F72L, and acylation in Q4. Our analysis indicates that the apparent change in rate-limiting step causes the observed epistatic effects. In other words, because substrate binding is slow in wtOXA-48, it is likely that neither S212A nor T213A substantially affect activity in the wild-type background and require F72L to accelerate substrate binding for their improvements to take effect fully.

### F72L and alanine mutations orthogonally tune the protein dynamics

To understand how evolution modulated the conformational dynamics of OXA-48, we determined the crystal structures of F72L, Q5, and Q5 covalently bound to CAZ (PDB IDs: 8PEA, 8PEB and 8PEC, Tab. S6). To extract dynamical information from these structures, we performed ensemble refinements for wtOXA-48 (PDB ID: 4S2P^28^), F72L, and Q5 (Fig. 3a). Firstly, the overall architecture of the active site in Q5, including the catalytic residues, was generally maintained (Fig. S6). Despite the introduction of S212A and T213A in the β5-β6 loop, the overall shape of this loop is preserved. As expected, the ensemble refinement confirmed that F72L increased the protein flexibility, primarily resulting in the Ω-loop adopting various alternate conformations. This increase in Ω-loop dynamics likely accelerated substrate binding and caused the observed change in rate-limiting step.^19, 29^

To understand how evolution affected the conformational dynamics while giving rise to synergy, we performed extensive molecular dynamics (MD) simulations of various OXA-48 variants in the acyl-enzyme complex (Fig. S7 to S11). We determined per-residue root-mean-square fluctuations (RMSF) for all variants to explore how evolution affected enzyme flexibility (Fig. 3b and S8). Consistent with the crystallographic observations, introducing F72L increased the RMSF values of the Ω loop region. While our simulations are of the acyl-enzyme state, the increased flexibility will likely also aid substrate entry.^19, 29^

To further dissect how the mutations affect the protein dynamics, we analyzed the conformational landscape of the active-site loops by principal component and cluster analysis (Fig. 3c, S9, and S10a). As expected, F72L significantly modulated the conformational landscape, while introducing the S212A or T213A mutations did not change the sampled conformational space (Fig. 3c and Fig. S10a). Notably, F72L allows the protein to populate a different conformational ensemble, in which the space created by the loss of the phenylalanine sidechain allows the neighboring α-helix to slide toward the center of the protein while increasing the conformational freedom of the Ω-loop (Fig. 3d and S10b). That movement, which provides space for substrate entry, is accompanied by tightening of an H-bonding network below the Ω-loop involving T71, Y144, and N169 (Fig. S10c).

While our MD analysis supports that F72L accelerates substrate binding and thereby shifts the rate-limiting step, the structural role of S212A and T213A is more elusive. Since these mutations did not affect the sampled conformational space, we hypothesized that dynamical cross-correlation analysis might allow us to identify regions that show correlated movements – and thus likely tighter interaction – with the substrate covalently bound to S70 (Fig. 3e and S11). When analyzing the changes in the correlation of the acylated S70 with the rest of the protein, we observed distinct dynamical changes wrought by either F72L or the alanine mutations on the protein scaffold. F72L primarily decreased the dynamical correlation between S70 and the Ω-loop, which agrees with the increased flexibility and improved substrate binding conferred by F72L. In contrast to F72L, S212A and T213 had an entirely different effect on the protein dynamics. Notably, S212A and T213A enhanced the correlation of S70 with the neighboring β-strands which harbor the oxyanion hole (Fig. 3e and S11).^30^ Dynamical correlations between the nucleophile and the oxyanion hole may explain increases in *k*_cat_, because they signify better preorganization and the ability to stabilize the transition state intermediate.^15, 31, 32^ Overall, the effects of F72L and the alanine mutations on the correlations were largely orthogonal, like their effects on conformational sampling. Such orthogonal dynamical relationships are probably central to the epistatic effects not only in OXA-48, but also for the evolution of other natural and designer enzymes.^25, 33^

**Figure 3:**
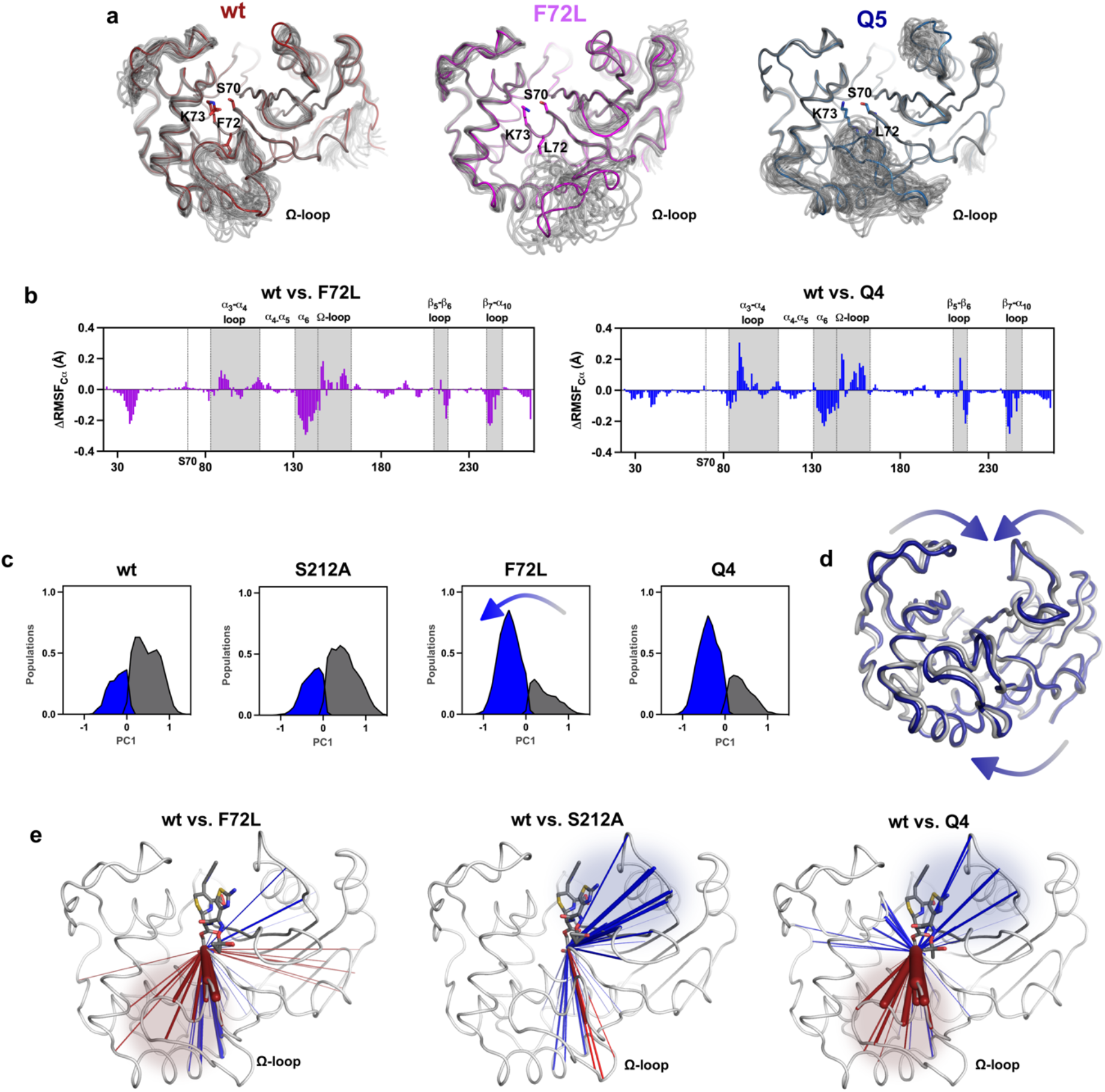
Evolution of a catalytically superior ensemble. **a.** Ensemble refinement of wtOXA-48, F72L, and Q5 reveal increased mobility of the Ω-loop. **b.** ΔRMSF values relative to wtOXA-48, F72L, and Q4 from MD simulations reproduce the increased flexibility of the Ω-loop region (see Fig. S8 for other variants). **c.** Principal component and cluster analysis show that F72L modulates the conformational landscape in ways likely to accelerate binding (see Fig. S9 for other variants). **d.** Cluster representatives indicate that evolution displaced the Ω-loop but closed the other solvent-exposed loops (see Fig. S10 for other variants). **e**. Dynamical correlation analysis reveals that the movement of the acylated S70 becomes tightly coupled with the protein scaffold, particularly the oxyanion hole, by means of the alanine mutations. In contrast, F72L predominantly decreases the interaction of S70 with the Ω-loop (increased and decreased correlations relative to wtOXA-48 are shown in blue and red, respectively. Only statistically significant changes compared to wtOXA-48 are shown (T-test, α = 0.05; see Fig. S11 for other variants).

## DISCUSSION

Unraveling the molecular mechanisms underlying epistasis is vital for understanding enzyme evolution.^1–4^ Many previous studies on epistatic enzyme evolution focused on exploring epistatic mechanisms from a structural perspective, for instance, by demonstrating direct interactions between mutations or synergistic effects on conformational dynamics.^7, 11, 34^ By contrast, our observations highlight that the molecular basis for intramolecular epistasis is rooted in changes in the catalytic cycle. We describe how distinct mutations shifted the rate-limiting step from substrate binding to the chemical reaction in the β-lactamase OXA-48, thereby causing strong phenotypic epistasis (Fig. 1 and 2). Intriguingly, the adaptive mutations introduced into OXA-48 are structurally orthogonal, but mechanistically epistatic: F72L induces dynamical and structural perturbations that are largely independent of the S212A and T213A, and *vice versa* (Fig. 3). Since CAZ binding is rate-limiting, F72L must be incorporated first to accelerate binding and unlock the effect of S212A and T213A on the chemical step. In addition to the observed change in the slowest step from binding to chemistry, shifting the rate-limiting step between other stages of the catalytic cycle could also result in epistasis. While genetic context-dependent effects on *k*_cat_ and *K*_M_ have been previously reported,^35^ deciphering their underlying epistatic relationship has remained challenging. Here, gaining detailed insights into the origins of epistasis was only possible by in-depth characterizations through pre-steady state kinetics, isotope effects, and dynamical analysis. To the best of our knowledge, this is the first report on how changing the reaction bottleneck from binding to the chemical step leads to synergy in evolution. We hypothesize that similar effects are likely to be of vast importance in the evolution of other biocatalysts.

’Catalytically perfect’ enzymes, in which the reaction rate is only limited by substrate diffusion, display catalytic efficiencies (*k*_cat_/*K*_M_) of >10^8^ M^-1^ s^-1^. Despite its *k*_cat_/*K*_M_ being orders of magnitude below the diffusion limit, our analysis unexpectedly revealed that CAZ binding is the rate-limiting step in OXA-48 (Fig. 2). In contrast, acylation and deacylation are often catalytic bottlenecks in serine β-lactamases.^21, 36^ Nevertheless, the extended size of CAZ has led to the hypothesis that its binding is more challenging than that of other β-lactams.^12, 36^ OXA-48 furthermore differs from other serine β-lactamases in that it accommodates the carboxylate group adjacent to the oxyimino-moiety of CAZ within its active site (Fig. S12),^20, 24^ which probably additionally slows down CAZ binding resulting in rate-limiting complex formation. Understanding how individual steps affect overall catalytic activity is crucial for understanding and predicting enzyme evolution and designing more efficient drugs that exploit mechanistic bottlenecks.

Our findings show the selection of a super-stoichiometric burst phase that correlates with increasing levels of antibiotic resistance (Fig. 2). The origin of such a burst phase cannot be described by a simple two-step reaction mechanism and is probably driven by the selection of pre-existing conformational sub-states and changes in conformational dynamics (Fig. 3).^11, 14, 37, 38^ The evolution of burst phase kinetics introduces a kinetic bottleneck that restricts efficiency at CAZ concentrations above *K*_M_ (Fig. 2b). Importantly, this bottleneck is irrelevant at the substrate concentrations used during selection, where the burst behavior vanishes. We hypothesize that offsetting activity enhancements under physiologically irrelevant steady-state conditions likely relieved selection pressure and facilitated improvements under the selective conditions.

In conclusion, our study demonstrates that mutations can orthogonally impact different steps along the catalytic cycle, ultimately shifting the catalytic bottleneck. The combined effect of these mutations results in positive epistasis, which drives enzyme evolution. Understanding the mechanistic origins of such epistatic phenomena is crucial to enhancing the predictability of evolutionary outcomes and advancing the overall field of enzyme engineering.

## Funding

The authors have declared no competing interest. CF thanks the PhD schools NFIF, IBA and Biocat for their funding. HAB thanks the SNSF for funding (P5R5PB_210999, PZ00P3_208691, P400PB_194329). MWvdK thanks BBSRC for funding (BB/M026280/1). NT thanks the Canadian Institute of Health Research (CIHR) for the project grant (AWD-019305). HKSL thanks CANS for the project grant. This work is part of a project that has received funding from the European Research Council under the European Horizon 2020 research and innovation program (PREDACTED Advanced Grant Agreement no. 101021207) to AJM. AJM and HAB also thank BBSRC (grant no. BB/R016445/1) and EPSRC (EP/M013219/1 and EP/M022609/1) for funding.

## Acknowledgements

Molecular dynamics simulations were conducted using the computational facilities of the Advanced Computing Research Centre, University of Bristol.

## MATERIALS AND METHODS

### General material

LB agar, broth, chloramphenicol, ampicillin, CAZ were purchased from Sigma-Aldrich (MO, USA). Primers (P) used for this study are shown in Tab. S7. All cloning enzymes were purchased from Thermo Fisher Scientific (MA, USA), if not stated otherwise. The *E. coli* E. cloni® 10G (MP21-5) was obtained from Lucigen (WI, USA). All strains used and constructed in this study are shown in Tab. S8. Kinetic data was fitted using Prism v. 9.0 (GraphPad Software, CA, USA).

### Directed evolution and cloning

The construction of the low copy number vector pUN-*bla*_OXA-48_ (pA15 origin; 10-20 copies per cell) was previously published.^39^ Error-prone PCR was performed using 10 ng pUNE-4-*bla*_OXA- 48_, GoTag (Promega, WI, USA), 25 mM MgCl_2_ (Promega), 10 µM P7/P8 and either 50 µM oxo-dGTP or 1 µM dPTP (Jena Bioscience, Germany). PCR products were *Dpn*I digested for 1 h and 37°C and 5 ng of each product were used for a second PCR, which was performed as described above, but without mutagenic nucleotides. PCR amplicons were digested using *Nco*I, *Xho*I, *Dpn*I for 1 h at 37°C, purified for ligation with the vector backbone and transformed into MP21-5. To insure a sufficient mutational depth, we aimed for library sizes of at least 5000 colonies which was determined by plating on agar plates supplemented with 25 mg/L chloramphenicol LB and one to two amino acid changes per round of evolution which was determined by Sanger sequencing (Azenta, Germany).

The fitness landscape was constructed in the pUN vector background using Goldengate cloning and the corresponding primers in Tab. S7. In short, we performed whole vector amplification followed by digestions with *Lgu*I and *DpnI* for 1 h 37°C. Ligations were performed using 10-20 ng of DNA for 1 h at room temperature using T4 ligase, transformed into MP21-5 and clones were grown on 25 mg/L chloramphenicol LB agar plates and verified using Sanger sequencing (Azenta).

For protein expression, OXA-48 variants were sub-cloned into a pDEST-17 (pURR) expression vector without the leader sequence and with a 6-His-tag using P2/P37 and P35/P36. Amplicons were digested using *Not*I and *Xho*I, ligated as described above and transformed into MP21-5. Vectors were isolated using the plasmid miniprep kit (Qiagen, Germany). pURR expression vectors were transformed into *E. coli* BL21 AI. Clones were selected on agar containing ampicillin 100 mg/L and verified using Sanger sequencing (Azenta).

### Selective plating

MP21-5 cultures harboring either pUNE-4-*bla*_OXA-48_ or a library of OXA-48 were plated on LB agar plates containing increasing concentrations of CAZ and grown over night at 37°C (Tab. S1). Up to eight colonies grown on the highest concentrations were recovered and their genotype characterized by Sanger sequencing (Azenta). Before determining their *IC_50_* values, the corresponding mutant alleles were sub-cloned into an isogenic pUN vector backbone and transformed into MP21-5, to exclude mutational effects outside of the target gene.

### *IC_50_* determination

*IC_50_* values were determined as described previously.^19, 39^ In brief, cultures were grown to full density under 700 rpm shaking over night at 37°C. Overnight cultures were diluted in PBS to a density of 10^6^ cells/ mL and used to inoculate a 384 well plate (Thermo Fisher Scientific) with a CAZ gradient (0 to 32 mg/L) at a final cell density of 10^5^ cells/ mL. Plates were incubated statically at 37°C for 20 h. The absorbance was determined as OD_600_ using an Epoch spectrophotometer (Biotek, VT, USA). Dose-response curves were and their *IC*_50_ value were determined based on a non-linear fit.

### Protein expression and purification

Cultures of *E. coli* BL21AI harboring modified pURR expression vector with *bla*_OXA-48_ or mutant alleles were grown in TB supplemented with 100 mg/L ampicillin at 30°C and 220 rpm. Protein expression was induced by adding L-arabinose (Sigma-Aldrich) to a final concentration of 0.2% when the cultures reached a OD_600_ of 0.4. Cultures were expressed for 16 h at 15°C, centrifuged at 4°C for 30 min and the cell pellets were stored at -20°C for purification. Protein purification was performed using HisPur^®^ Ni-NTA spin columns (Thermo Fisher Scientific) as published previously.^19, 40^ For crystallization, the His-tag was cleaved over night at 4°C using in-house produced TEV and the TEV-cleaved product was purified through an additional round of HisPur^®^ Ni-NTA columns (Thermo Fisher Scientific).

### Fluorescence-based thermostability

Thermostability was performed as previously published using purified OXA-48 enzymes, containing 6-His tag and TEV cleaving site.^19^ In 50 mM HEPES (VWR, PA, USA), pH 7.5 including 50 mM potassium sulfate (Honeywell, NC, USA), enzymes were diluted to 0.2 mg/mL and mixed with 5xSYPRO orange (Sigma-Aldrich). Using an MJ minicycler (Bio-Rad, CA, USA), a temperature gradient (25 to 70°C) was performed with a heating rate of 1°C per min was. All experiments were performed in triplicates and the melting temperatures (T_M_) were determined as the inflection point of the melting transition found from the first derivative.

### Pre-steady-state burst and steady-state enzyme kinetics

Room-temperature burst kinetics were obtained under pre-steady state conditions using an SX20 stopped flow (Applied Photophysics, UK) by monitoring substrate depletion by absorbance at 260 nm. Enzyme and substrate were mixed 1:1 at 25°C and in 0.1 M phosphate buffer (Sigma-Aldrich, pH 7.2) supplemented with 50 mM NaHCO_3_ (Sigma-Aldrich). Burst kinetics were assayed at final enzyme concentrations of 10 μM and final substrate concentrations varying between 50 to 400 μM (Eq. 1).

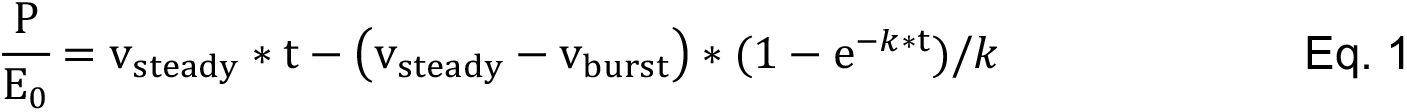

Catalytic parameters (*k*_cat_*, K*_M_ and *k*_cat_/*K*_M_) were determined under burst and steady-state conditions using ceftazidime (Δξ = −9,000 M^−1^ cm^−1^) at 260 nm by measuring the initial enzymatic reaction rate in a Epoch plate-reader (Biotek). Burst phase rates were determined at 4°C and steady-state parameters were determined at 25°C. Reactions rates were obtained in at least duplicates at a final enzyme concentration of 1 μM (final assay volume of 100 μL). UV-transparent 96-well plates (Corning, ME, USA) were used. Assays were performed in 0.1 M phosphate buffer (Sigma-Aldrich, pH 7.2), supplemented with 50 mM NaHCO_3_ (Sigma-Aldrich).

### Solvent isotope effects

Solvent isotope effects were determined at 25°C using an SX20 stopped flow (Applied Photophysics) from burst kinetics obtained at 400 μM CAZ. Solvent isotope effects (KIE) were calculated from the ratio of the rate in 80% D_2_O (Sigma-Aldrich, *k*_D_) and water (*k*_H_, Eq. 2).

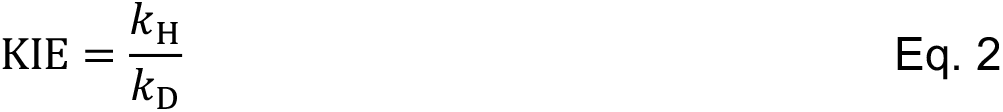

### Pre-steady state binding kinetics

CAZ binding was determined under pre-steady state condition using an SX20 stopped flow (Applied Photophysics) by tryptophane fluorescence, with an excitation wavelength of 280 nm, a 305 nm lower cut off emission filter and a 0.2 mm excitation pathlength. Enzyme and substrate were mixed 1:1 at a final enzyme concentration of 1 μM, and the final substrate concentration was varied between 100 to 1200 μM. Binding was assayed at 25°C and in 0.1 M phosphate buffer (pH 7.2) supplemented with 50 mM NaHCO_3_ (Sigma Aldrich). Binding rates were obtained by global fitting of the observed biphasic curves to a double-exponential decay (Eq. 3).

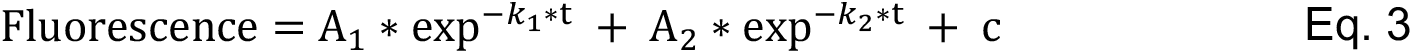

where *k*_1_ and *k*_2_ are the observed binding rates and A_1_ and A_2_ are the amplitudes of the two signals with an offset of c. To enable global fitting, *k*_1_ and *k*_2_ were fitted to a linear equation each (Eq. 4, Eq. 5), where *k*_1,on_, *k*_2,on_, *k*_1,off_ and *k*_2,off_ were shared between all datasets.

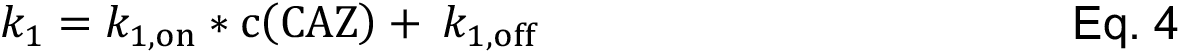

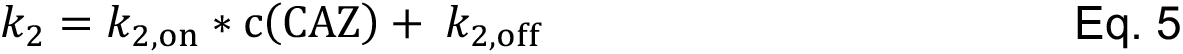

### Size exclusion chromatography

Changes in the molecular size were understudied using size exclusion chromatography: 50 nM, 10 μM wtOXA-48 and 10 μM of the mutant R189A/R206A mutants. R189A/R206A has been previously described to disrupt the dimer interface of OXA-48 and elute significantly later.^41^ Separation was performed in 0.1 M phosphate buffer (Sigma-Aldrich, pH 7.0) using a Superdex 200 10/300 GL column with a flow rate of 0.5 mL per min at 4°C. Elution was monitored by recording the absorbance at 280 nm.

### Dynamic light scattering

The hydrodynamic radius of Q4 was determined using a zetasizer (Malvern Panalytical, UK). To that end, 1 mL of 10 μM Q4 was assayed either in the presence or absence of 400 μM CAZ in 0.1 M phosphate buffer (Sigma-Aldrich, pH 7.0) supplemented with 50 mM NaHCO_3_ (Sigma-Aldrich). The experiment with CAZ was repeated after a 15 min incubation.

### Sequential mixing

To assess the reversibility of the burst phase, 20 μM Q4 was pre-incubated with 400 μM CAZ, and mixed with a second batch of 400 μM CAZ after a delay of 1000, 2000, and 3000 s in an SX20 stopped flow (Applied Photophysics) by monitoring substrate depletion by absorbance at 260 nm.

### Crystallography, structure determination and refinement

Crystallization was performed in a 1 μL hanging drop containing 10 mg/mL enzyme and mixed 1:1 with reservoir solution containing 0.1 M Tris, pH 9.0 (Sigma-Aldrich), and 28 to 30% polyethylene glycol (PEG) mono ethylene ether 500 (Sigma-Aldrich) at 4°C. Crystals were cryoprotected by using 15% ethylene glycol (Sigma-Aldrich) in addition to the reservoir solution, and subsequently frozen in liquid nitrogen. Diffraction data were collected on ID23- EH2 (F72L) and ID30B (Q5 and Q5-CAZ), ESRF, France, at 100 K, wavelength 0.9184 Å, and the diffraction images were indexed and integrated using XDS^42^. For data scaling, AIMLESS was used^43^ and an overall high completeness and CC1/2 > 0.5 and a mean intensity above 1.0 in the outer resolution shell was aimed for (Tab. S6). Molecular replacement was performed using chain A of PDB ID 6Q5F^19^ and the program PHENIX 1.12^44^. Parts of the model were manually rebuilt using Coot^45^. Average structure refinement and ensemble refinement was performed using PHENIX 1.12. PyMOL 1.8 was used for illustrations (Schrödinger, NY, USA).

### Molecular dynamics simulations: System setup

MD simulations were set up analogously to our previous work.^19, 24^ See Fig. S7 to S11 for all simulated structures. Acyl-enzyme structures of the OXA-48 variants with covalently bound CAZ were built based on the structure of apo wtOXA-48 (PDB ID: 4S2P^28^), with CAZ added from the holo structure of Q5 (PDB ID: 8PEC). All OXA-48 variants were modelled based on this structure using the mutagenesis tool in PyMOL, choosing the rotamer with the least steric clashes with surrounding atoms. For comparison, MD simulations for Q4 were also performed based on the holo structure of Q5 (PDB ID: 8PEC) and the Ω-loop of apo wtOXA-48 (residues D143 – I164). This variant is indicated as Q4 (template Q5) in Fig. S7 - S11. The results from those simulations agreed qualitatively well with those obtained from the Q4. The system was parametrized using tleap,^46^ and enzymes were solvated in a 10.0 Å octahedral box of TIP3P water^47, 48^ with net charge neutralized using the Amber uniform neutralizing plasma.^46^ The ff14SB force field^49^ was used to describe the protein. Parameters for the carbamylated lysine (KCX) were previously obtained^24^ from restrained electrostatic potential (RESP) fitting as implemented in the RED Server.^50^ Parameters for the CAZ-acetylated serine were likewise obtained with the RED Server.

Several restraints were applied during the simulations to maintain a productive conformation. The restraints include a ≤ 4.0 Å distance restraint from the nucleophilic water to either the KCX base oxygen or CAZ carbonyl carbon using one-sided harmonic potentials. The distance of the KCX N_ζ_ and catalytic Ser70 O_γ_ was likewise restrained to ≤ 4.0 Å. Lastly, flat-bottom potentials were applied to the C_δ_C_ε_N_ζ_C_η_ (≤ -130° and ≥ -80°) and C_γ_C_δ_C_ε_N_ζ_ (≤ 45° and ≥ 5=95°) dihedral angles. All restraint force constants were 10 kcal/mol/Å^2^ during the equilibration MD and 100 kcal/mol/Å^2^ during minimization.

### Molecular dynamics simulations: Simulations

All systems were minimized using 10,000 steps of steepest descent followed by 10,000 steps of conjugate gradient. During both minimization steps, the position of all protein atoms was restraint with a weight of 10 kcal/mol/Å^2^. The minimization was subsequently repeated without positional restraints. Subsequently, the system was heated from 50 K to 300 K in 20 ps, and then simulated for 50 ns in the NPT ensemble saving a frame every 100 ps. Langevin dynamics were used with a collision frequency of 0.2 and a 2 fs time step. The Berendsen barostat was used with isotropic position scaling. All bonds involving hydrogens were constrained using the SHAKE algorithm. 20 independent simulations were run per enzyme variant (for a total of 1.0 µs per variant). All calculations were performed with the Amber18 program package (sander.MPI for minimization and pmemd.cuda for MD simulations).^46^

### Molecular dynamics simulations: Analysis

MD simulations were analyzed using CPPTRAJ^51^. All analyses were based on C_α_ positions. The first 10 ns of each production MD run were excluded to allow sufficient time for system equilibration. RMSD values were calculated compared to the minimized starting structures. Root mean square fluctuations (RMSF) were determined by first calculating an average structure for each replicate, aligning the trajectory against the average structure, and then calculating the RMSF for each protein residue. Errors indicate the standard error of the 20 independent replicates.

For each variant, cluster and principal component analysis were performed on their combined 20 replicates. To that end, a global average structure over all variants was first determined and all trajectories snapshots were aligned by Cα to that average. After the alignment, the analyses were performed for the active-site loops (residues 96-106, 151-160, 213-218, 242- 246) without re-aligning the loops. To perform the analyses in the same space, both the cluster and principal component analysis were performed simultaneously for all variants. Clustering based on Cα RMSD of the loops was performed with cpptraj using the kmeans algorithm to split each trajectory into two clusters. Principal component analysis was performed using mdtraj^52^ and sklearn.^53^

