## Supplementary information for "Epistasis Arises from Shifting the Rate-Limiting Step during Enzyme Evolution"

**Contents**

1. Supplementary tables 2
Table S1: $IC_{50}$ values against ceftazidime for variants selected during directed evolution. 2
Table S2: Overview of $IC_{50}$ values for all mutational combinations of Q4. 2
Table S3: Test for significant changes in $IC_{50}$ for single mutants. 3
Table S4: Melting temperatures of OXA-48 variants. 3
Table S5: Steady state kinetics determined at 25ºC. 3
Table S6: X-ray data collection and standard refinement statistics. 4
Table S7: Primers used in this study. 5
Table S8. List of strains used and constructed in this study. 6
2. Supplementary figures 7
Figure S1: Epistasis across the fitness landscape. 7
Figure S2: Burst-phase kinetics. 8
Figure S3: Steady-state Michaelis-Menten kinetics. 9
Figure S4: Correlation of *in vitro* $k_{cat}/K_M$ values with *in vivo* $IC_{50}$ values. 10
Figure S5: Sequential mixing (burst recovery) and size exclusion chromatography. 11
Figure S6: Conserved active site architecture between wtOXA-48 and Q5. 12
Figure S7: MD simulations of the OXA-48 variants. 13
Figure S8: Changes in per-residue $C_\alpha$ RMSF values compared to wtOXA-48. 14
Figure S9: Principal component (PC) analysis of the OXA-48 variants analysis of active-site loops (residues 96-106, 151-160, 213-218, 242-246). 15
Figure S10: Cluster analysis of the OXA-48 variants. 16
Figure S11: Dynamical correlations in the OXA-48 variants. 17
Figure S12: Ceftazidime orientation in different β-lactamases. 18
3. References 19

### 1. Supplementary tables

**Table S1: $IC_{50}$ values against ceftazidime for variants selected during directed evolution.**

| Strain no.[a] | Round of evolution | Ceftazidime selection concentration (mg/L) | Mutational background | Ceftazidime $IC_{50}$ (mg/L) |
|---|---|---|---|---|
| MP21-5 | - | - | - | 0.012 |
| MP21-1 | - | - | wild-type OXA-48 | 0.013 |
| MP24-51 | Round 1 | 0.25 | F72L | 0.029 |
| MP24-52 | Round 2 | 2 | F72L/S212A | 0.140 |
| MP24-53 | Round 3 | 2 | F72L/S212A/T213A (Q3) | 0.389 |
| MP24-54 | Round 4 | 8 | A33V/F72L/S212A/T213A (Q4) | 0.513 |
| MP24-55 | Round 5 | 4 | A33V/K51E/F72L/S212A/T213A (Q5) | 0.556 |

[a] see Tab. S8 for a full description of all strains used in this work

**Table S2: Overview of $IC_{50}$ values for all mutational combinations of Q4.**

| Strain no.[a] | Amino acids[b] | Number of mutations[c] | Ceftazidime $IC_{50}$ (mg/L) |
|---|---|---|---|
| MP21-1 | AFST | 0 | 0.013 ± 0.002 |
| MP22-7 | AFSA | 1 | 0.012 ± 0.001 |
| MP22-6 | AFAT | 1 | 0.015 ± 0.002 |
| MP22-8 | AFAA | 2 | 0.023 ± 0.005 |
| MP22-5 | ALST | 1 | 0.029 ± 0.006 |
| MP22-38 | ALSA | 2 | 0.177 ± 0.063 |
| MP22-37 | ALAT | 2 | 0.140 ± 0.003 |
| MP22-39 | ALAA (Q3) | 3 | 0.389 ± 0.025 |
| MP22-20 | VFST | 1 | 0.017 ± 0.004 |
| MP22-42 | VFSA | 2 | 0.015 ± 0.001 |
| MP22-41 | VFAT | 2 | 0.014 ± 0.001 |
| MP12-47 | VFAA | 3 | 0.020 ± 0.003 |
| MP22-40 | VLST | 2 | 0.034 ± 0.002 |
| MP12-46 | VLSA | 3 | 0.141 ± 0.028 |
| MP12-45 | VLAT | 3 | 0.148 ± 0.042 |
| MP12-48 | VLAA (Q4) | 4 | 0.513 ± 0.093 |

[a] see Tab. S8 for a full description of all strains used in this work.

[b] amino acid code referring to the A33V, F72L, S212A and T213A

[c] compared to wtOXA-48.

**Table S3: Test for significant changes in $IC_{50}$ for single mutants.**

| Test details[a] | Adjusted *P* value |
|---|---|
| wtOXA-48 vs. A33V | 0.7048 |
| wtOXA-48 vs. F72L | 0.0060 |
| wtOXA-48 vs. S212A | 0.1113 |
| wtOXA-48 vs. T213A | 0.9815 |

[a] Brown-Forsythe one way ANOVA (df=4, $P < 0.001$) followed by Dunnett post hoc test with wtOXA-48 as a control group

**Table S4: Melting temperatures of OXA-48 variants.**

| OXA-48 variants | Melting temperatures (°C) |
|---|---|
| wild-type | 52.9 ± 0.1 |
| A33V | 52.7 ± 0.1 |
| F72L | 45.7 ± 0.1 |
| S212A | 52.5 ± 0.1 |
| T213A | 53.3 ± 0.1 |
| F72L/S212A | 45.8 ± 0.1 |
| F72L/T213A | 45.6 ± 0.1 |
| F72L/S212A/T213A (Q3) | 45.0 ± 0.1 |
| A33V/F72L/S212A/T213A (Q4) | 45.3 ± 0.1 |
| A33V/K51E/F72L/S212A/T213A (Q5) | 44.2 ± 0.1 |

Errors are reported as the standard deviation.

**Table S5: Steady state kinetics determined at 25ºC.**

| | $k_{cat}$ ($s^{-1}$) | $K_M$ (μM) | $k_{cat}$ / $K_M$ ($M^{-1}$ $s^{-1}$) |
|---|---|---|---|
| wtOXA-48 | 0.0028 ± 0.0008 | 247 ± 140 | 11 |
| F72L | 0.0049 ± 0.0004 | 18 ± 5 | 281 |
| S212A | NC[a] | NC[a] | 24 |
| T213A | NC[a] | NC[a] | 31 |
| F72L/S212A | 0.0037 ± 0.0003 | 7 ± 3 | 565 |
| F72L/T213A | 0.0180 ± 0.0005 | 53 ± 5 | 339 |
| S212A/T213A | NC[a] | NC[a] | 98 |
| Q3 | 0.0068 ± 0.0003 | 2 ± 1 | 3222 |
| Q4 | 0.0101 ± 0.0004 | 7 ± 2 | 1555 |

[a] NC: Not calculatable due to linearity of the Michaelis-Menten plot.
Errors are reported as the standard deviation of the mean.

**Table S6: X-ray data collection and standard refinement statistics.**

| | F72L | Q5 | Q5-CAZ |
|---|---|---|---|
| PBP | 8PEA | 8PEB | 8PEC |
| Beamline | ID23-EH2, ESRF | ID30B, ESRF | ID30B, ESRF |
| Wavelength (Å) | 0.8731 | 0.9763 | 0.9763 |
| Resolution range (Å) | 41.41-1.97 (2.04-1.97) | 24.95-1.17 (1.19-1.17) | 24.39-2.66 (2.78-2.66) |
| Space group | P $2_12_12_1$ | C2 | P $6_1$ |
| Unit cell: a,b,c (Å) | 64.68, 82.82, 100.99 | 94.40, 42.54, 64.30 | 202.15, 202.15, 55.70 |
| α, β, γ (°) | 90, 90, 90 | 90, 106.88, 90 | 90, 90, 120 |
| Total reflections | 277469 (24690) | 356104 (16085) | 230393 (28723) |
| Unique reflections | 38991 (3822) | 82172 (4063) | 37514 (4578) |
| Multiplicity | 7.1 (6.5) | 4.3 (4.0) | 6.1 (6.3) |
| Completeness (%) | 99.74 (99.74) | 99.7 (99.9) | 99.2 (100.0) |
| Mean I/sigma(I) | 10.05 (0.99) | 8.7 (1.1) | 9.1 (1.0) |
| Overall *B*-factor from Wilson plot (Å$^2$) | 37.28 | 12.7 | 73.12 |
| $R_{merge}$ | 0.1072 (1.352) | 0.067 (0.798) | 0.108 (1.555) |
| $R_{measured}$ | 0.1159 (1.474) | 0.086 (1.244) | 0.129 (1.846) |
| $R_{pim}$ | 0.04355 (0.5769) | 0.041 (0.615) | 0.052 (0.761) |
| $CC_{1/2}$ | 0.998 (0.675) | 0.996 (0.521) | 0.997 (0.367) |
| Resolution range (Å) | 41.41-1.97 | 24.95-1.17 | 24.39-2.66 |
| Reflections used in refinement | 38958 (3818) | 82157 (8180) | 37109 (3390) |
| Reflections used for R-free | 1613 (159) | 1248 (113) | 1809 (181) |
| Final $R_{work}$ | 0.2045 (0.3786) | 0.1647 (0.3070) | 0.2028 (0.3291) |
| Final $R_{free}$ | 0.2474 (0.3992) | 0.1871 (0.2873) | 0.2722 (0.3682) |
| No. of non-hydrogen atoms | 4264 | 2557 | 7669 |
| -macromolecules | 3968 | 2183 | 7472 |
| -ligands | 1 | 22 | 113 |
| -solvent | 295 | 352 | 84 |
| R.m.s. deviations | | | |
| -bonds (Å) | 0.009 | 0.009 | 0.009 |
| -angles (°) | 1.19 | 1.16 | 1.09 |
| Ramachandran plot | | | |
| -Favoured (%) | 94.58 | 98.75 | 92.52 |
| -Allowed (%) | 3.33 | 1.25 | 7.03 |
| -Outliers (%) | 0.95 | 0.00 | 0.45 |
| Average B-factor (Å$^2$) | 45.74 | 23.67 | 96.35 |
| -macromolecules (Å$^2$) | 45.69 | 21.77 | 96.57 |
| -ligands (Å$^2$) | 35.79 | 27.91 | 94.36 |
| -solvent (Å$^2$) | 46.41 | 35.18 | 78.99 |

Statistics for the highest-resolution shell are shown in parentheses.

**Table S7: Primers used in this study.**

| No. | Name | | 5´-3´ | Ref. |
|---|---|---|---|---|
| 7 | preOXAseq | F | GATTACGCGCAGACCAAAACG | [1] |
| 8 | postOXAseq | R | CCTATTTCCCTAAAGGGTTTATTGAGAATATG | [1] |
| 15 | F72L | F | TTTTTTGCTCTTCGCATCTACCCTGAAAATTCCCAATAGCTT | [1] |
| | | R | TTTTTTGCTCTTCGATGCGGGTAAAAATGCTTG | |
| 16 | S212A | F | TTTTTTGCTCTTCACTGGATACGCGACTAGAATCGAACCTAAGATTGG | [1] |
| | | R | TTTTTTGCTCTTCCCAGTTTTAGCCCGAATAATATAGTCACC | |
| 17 | T213A | F | TTTTTTGCTCTTCACTGGATACTCGGCGAGAATCGAACCTAAGATTGG | [1] |
| | | R | P16R | |
| 18 | S212A/T213A | F | TTTTTTGCTCTTCACTGGATACGCGGCGAGAATCGAACCTAAGATTGG | This study |
| | | R | P16R | |
| 25 | A33V | F | TTTTTTGCTCTTCAGTTGGAATGTTCACTTTACTGAACAT | This study |
| | | R | TTTTTTGCTCTTCCAACTTTTGTTTTCTTGCCATTC | |
| 35 | pDest17 vector-NotI | F | TTTTTTGCGGCCGCTTCGAGGTGATGGTGATGGTGATGGTAGTACGACATA | This study |
| 36 | pDest17 vector-XhoI | R | TTTTTTCTCGAGTGATTCGAGGCTGCTAACAAAGCCCG | This study |
| 37 | NotI-TEV-OXA-48 | F | TTTTTTGCGGCCGCAGAGAACCTGTATTTTCAGGGTAAGGAATGGCAAGAAAACAAAAGTTGG | This study |
| | | R | P2R | |

F: forward primer, R: reverse primer

**Table S8. List of strains used and constructed in this study.**

| Strain no. | Strain name [a] | Vector number | Insert | Reference |
|---|---|---|---|---|
| 21-5 | E cloni | None | None | Lucigen |
| 21-1 | E cloni | pUNe-4 | wtOXA-48 | [1] |
| Clones selected after error prone PCR: | | | | |
| 21-9 | E cloni | pUNe-5 | F72L | This study |
| 21-21 | E cloni | pUNe-8 | F72L/S212A | This study |
| 21-30 | E cloni | pUNe-11 | F72L/S212A/T213A | This study |
| 21-62 | E cloni | pUNe-14 | A33V/F72L/S212A/T213A | This study |
| 21-73 | E cloni | pUNe-17 | A33V/K51E/F72L/S212A/T213A | This study |
| Subcloned into isogenic vector and strain backbone after selection: | | | | |
| 24-51 | E.cloni | pUNe-5.1 | F72L | This study |
| 24-52 | E.cloni | pUNe-8.1 | F72L/S212A | This study |
| 24-53 | E.cloni | pUNe-11.1 | F72L/S212A/T213A | This study |
| 24-54 | E.cloni | pUNe-14.1 | A33V/F72L/S212A/T213A | This study |
| 24-55 | E.cloni | pUNe-17.1 | A33V/K51E/F72L/S212A/T213A | This study |
| Mutants constructed for the landscape: | | | | |
| 22-5 | E cloni | pUNs-1 | F72L | [1] |
| 22-6 | E cloni | pUNs-2 | S212A | [1] |
| 22-7 | E cloni | pUNs-3 | T213A | [1] |
| 22-8 | E cloni | pUNs-4 | S212A/T213A | This study |
| 22-20 | E cloni | pUNs-17 | A33V | This study |
| 22-37 | E cloni | pUNs-34 | F72L/S212A | [1] |
| 22-38 | E cloni | pUNs-35 | F72L/T213A | This study |
| 22-39 | E cloni | pUNs-36 | F72L/S212A/T213A | This study |
| 22-40 | E cloni | pUNs-37 | A33V/F72L | This study |
| 22-41 | E cloni | pUNs-38 | A33V/S212A | This study |
| 22-42 | E cloni | pUNs-39 | A33V/T213A | This study |
| 12-45 | E cloni | pUNs-121 | A33V/F72L/S212A | This study |
| 12-46 | E cloni | pUNs-122 | A33V/F72L/T213A | This study |
| 12-47 | E cloni | pUNs-123 | A33V/S212A/T213A | This study |
| 12-48 | E cloni | pUNs-124 | A33V/F72L/S212A/T213A | This study |
| Mutants constructed for protein expression (without signal peptide): | | | | |
| 12-71 | E cloni | pURR-1 | wtOXA-48 | This study |
| 12-72 | E cloni | pURR-2 | F72L | This study |
| 12-73 | E cloni | pURR-3 | S212A | This study |
| 12-74 | E cloni | pURR-4 | T213A | This study |
| 12-75 | E cloni | pURR-5 | F72L/S212A | This study |
| 12-76 | E cloni | pURR-6 | F72L/T213A | This study |
| 12-77 | E cloni | pURR-7 | F72L//S212A/T213A | This study |
| 24-16 | E.cloni | pURR-14 | A33V/F72L/S212A/T213A | This study |
| 13-78 | E.cloni | pDEST-17 | A33V/K51E/F72L/S212A/T213A | This study |
| Strains for protein expression: | | | | |
| 13-02 | BL21 AI | None | None | ThermoFisher |
| 24-01 | BL21 AI | pURR1 | wtOXA-48 | This study |
| 24-02 | BL21 AI | pURR-2 | F72L | This study |
| 24-03 | BL21 AI | pURR-3 | S212A | This study |
| 24-04 | BL21 AI | pURR-4 | T213A | This study |
| 24-05 | BL21 AI | pURR-5 | F72L/S212A | This study |
| 24-06 | BL21 AI | pURR-6 | F72L/T213A | This study |
| 24-07 | BL21 AI | pURR-7 | F72L//S212A/T213A | This study |
| 24-08 | BL21 AI | pURR-14 | A33V/F72L/S212A/T213A | This study |
| 13-80 | BL21 AI | pURR-11 | A33V/K51E/F72L/S212A/T213A | This study |

[a] Species for all strains is *E. coli.*

### 2. Supplementary figures

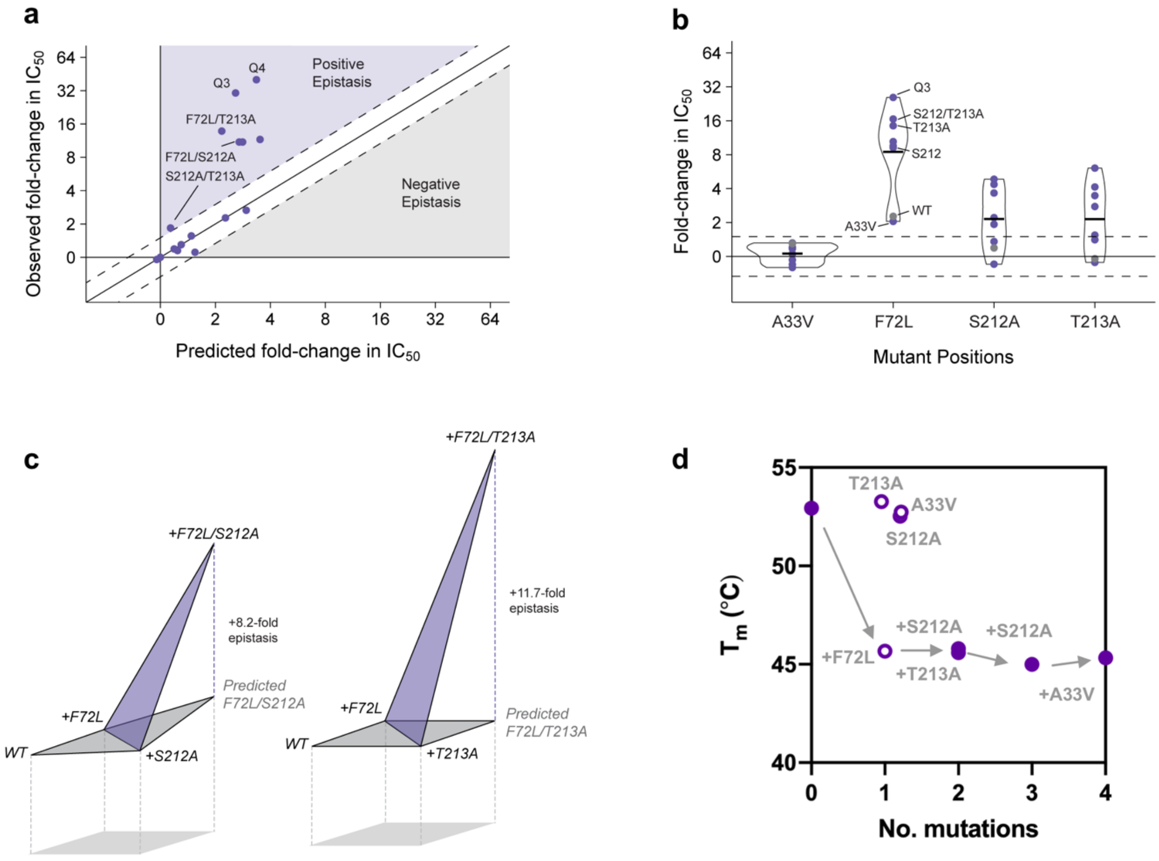


**Figure S1: Epistasis across the fitness landscape.**

**a.** Predicted $IC_{50}$ fold changes based on additive contributions of single mutational effects in wtOXA-48. Genotypes were positively epistatic (purple) when observed fold-changes exceeded additive predictions by more than 1.5-fold and negatively epistatic (grey) when less than 1.5-fold. **b.** Contribution of mutations on the $IC_{50}$ fold-change in each genotypic background. Distributions are shown by violin plots with bars representing mean fold-change in $IC_{50}$ and grey points representing the wtOXA-48 background. **c.** Positive epistasis is particularly apparent in the combinations of F72L with either S212A or T213A. **d.** Changes in melting temperature ($T_m$) for single and multistep mutants during the directed evolution. Open circles represent single mutants, and filled circles mutational combinations.

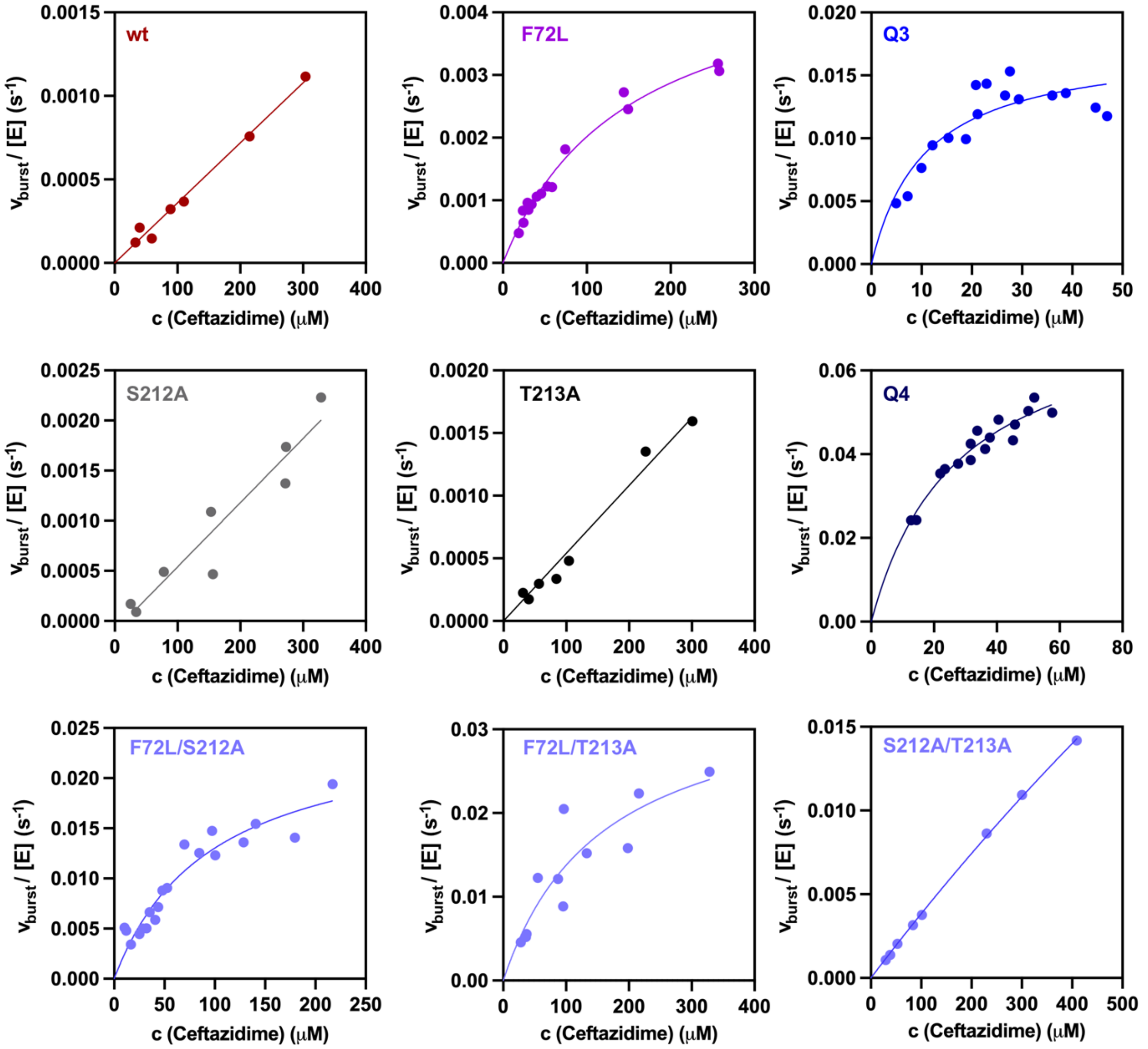


**Figure S2: Burst-phase kinetics.**

The kinetics reveal that F72L predominantly affects affinity ($K_M$), while S212A and T213A improve catalysis ($k_{cat}$). The *in vitro* burst-phase $k_{cat}/K_M$ values correlate well with the *in vivo* $IC_{50}$ values (Fig. S4). Burst-phase activities were determined at 4ºC.

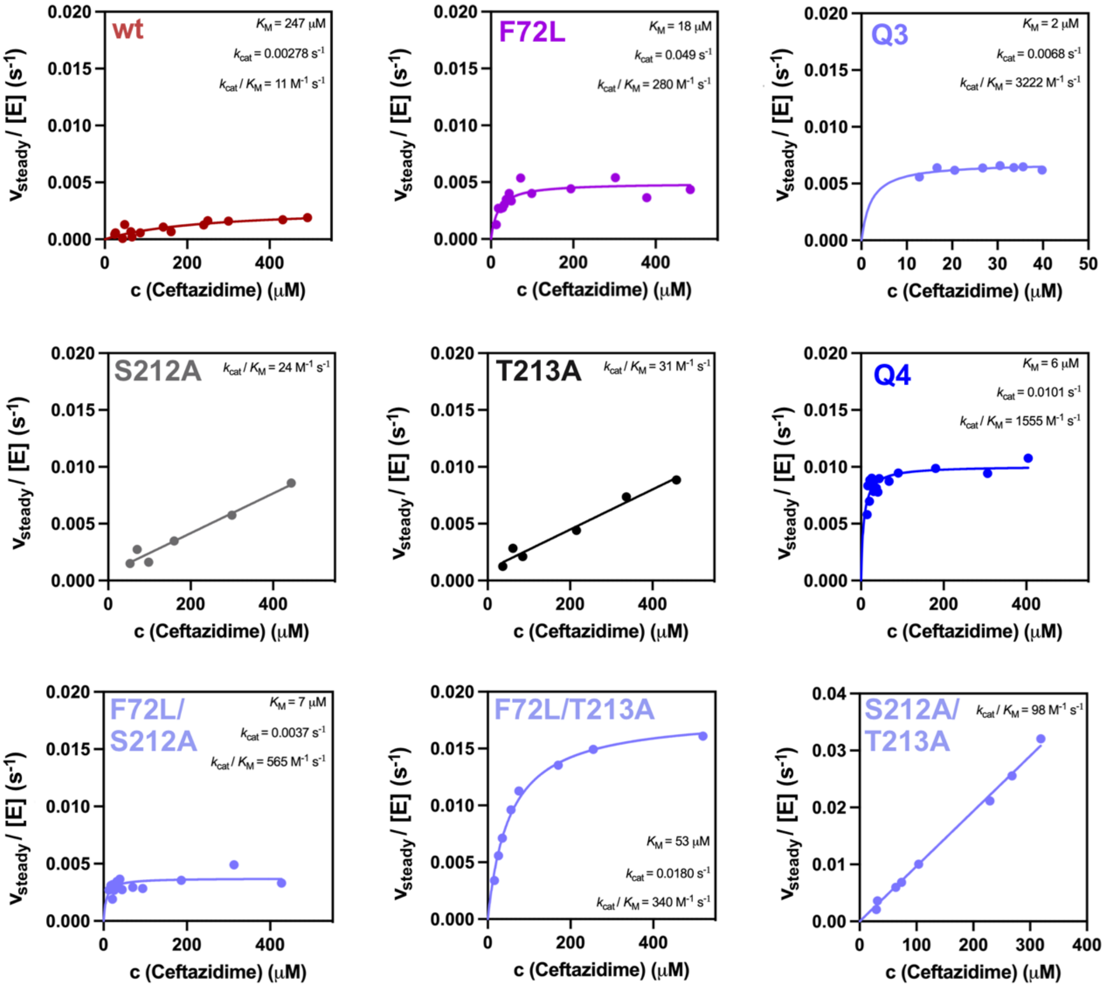


**Figure S3: Steady-state Michaelis-Menten kinetics.**

The Michaelis-Menten kinetics of the steady-state for various OXA-48 variants reveal similar overall trends compared to our burst phase results (Fig. S2). F72L predominantly lowers affinity ($K_M$), while S212A and T213A improve catalysis ($k_{cat}$). Nevertheless, the epistasis evident from the $IC_{50}$ values is not as pronounced in the steady-state as in the burst-phase kinetics (Fig. S2 and S4). Activities were determined at 25ºC.

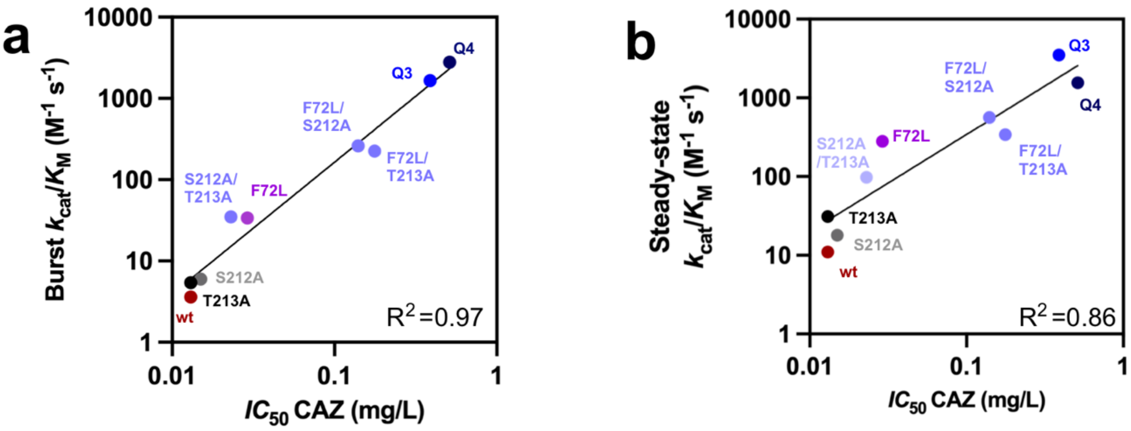


**Figure S4: Correlation of *in vitro* $k_{cat}/K_M$ values with *in vivo* $IC_{50}$ values.**

**a.** The burst-phases $k_{cat}/K_M$ values correlate mores strongly with $IC_{50}$ ($R^2$ = 0.97) than **b.** $k_{cat}/K_M$ values obtained from steady-state kinetics ($R^2$ = 0.86). Burst-phase data was obtained at 4ºC, and steady-state measurements were performed at 25ºC.

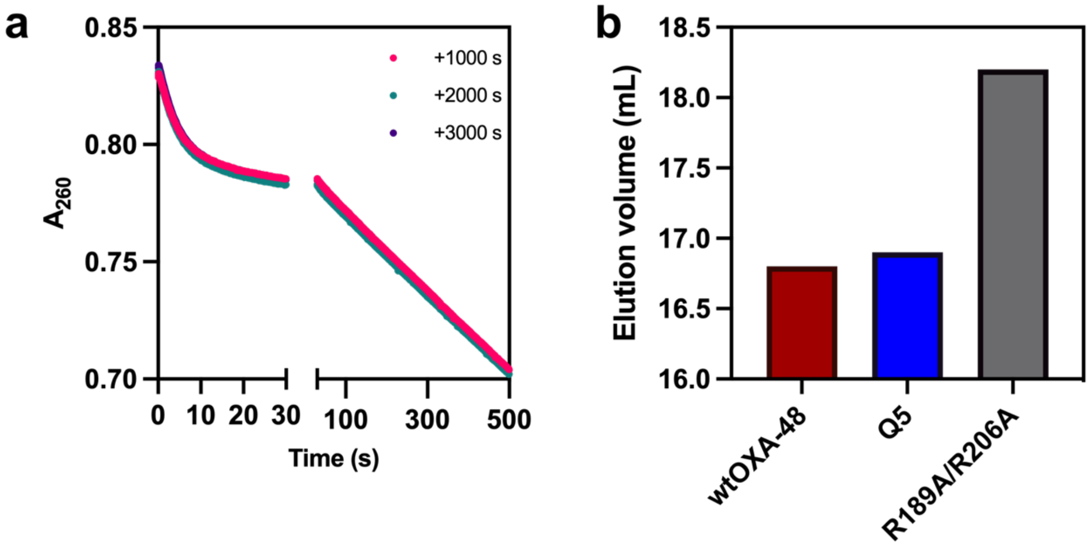


**Figure S5: Sequential mixing (burst recovery) and size exclusion chromatography.** **a.** Repeated sequential mixing using Q4 demonstrates the full recovery of the enzyme after each burst. **b.** Elution volume between wtOXA-48, Q5, and the monomeric wtOXA-48 mutant R189A/R206A[2] indicate that Q5 remains dimeric under the assay conditions.

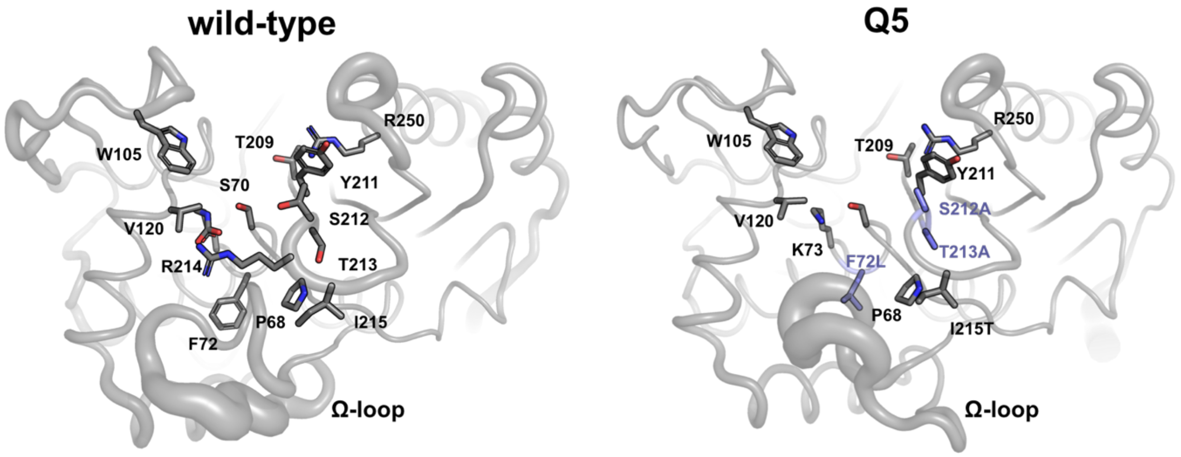


**Figure S6: Conserved active site architecture between wtOXA-48 and Q5.**

Relevant mutational sites, such as F72L, S212A and T213A, are colored in purple (Q5). Active site residues important for substrate binding and catalysis are displayed in both wtOXA-48 and Q5. The average refined crystal structure of Q5 displays a significant shift in the position of the Ω-loop. Ensemble refinements demonstrate a substantial increase in flexibility of the Ω-loop (Fig. 3a)

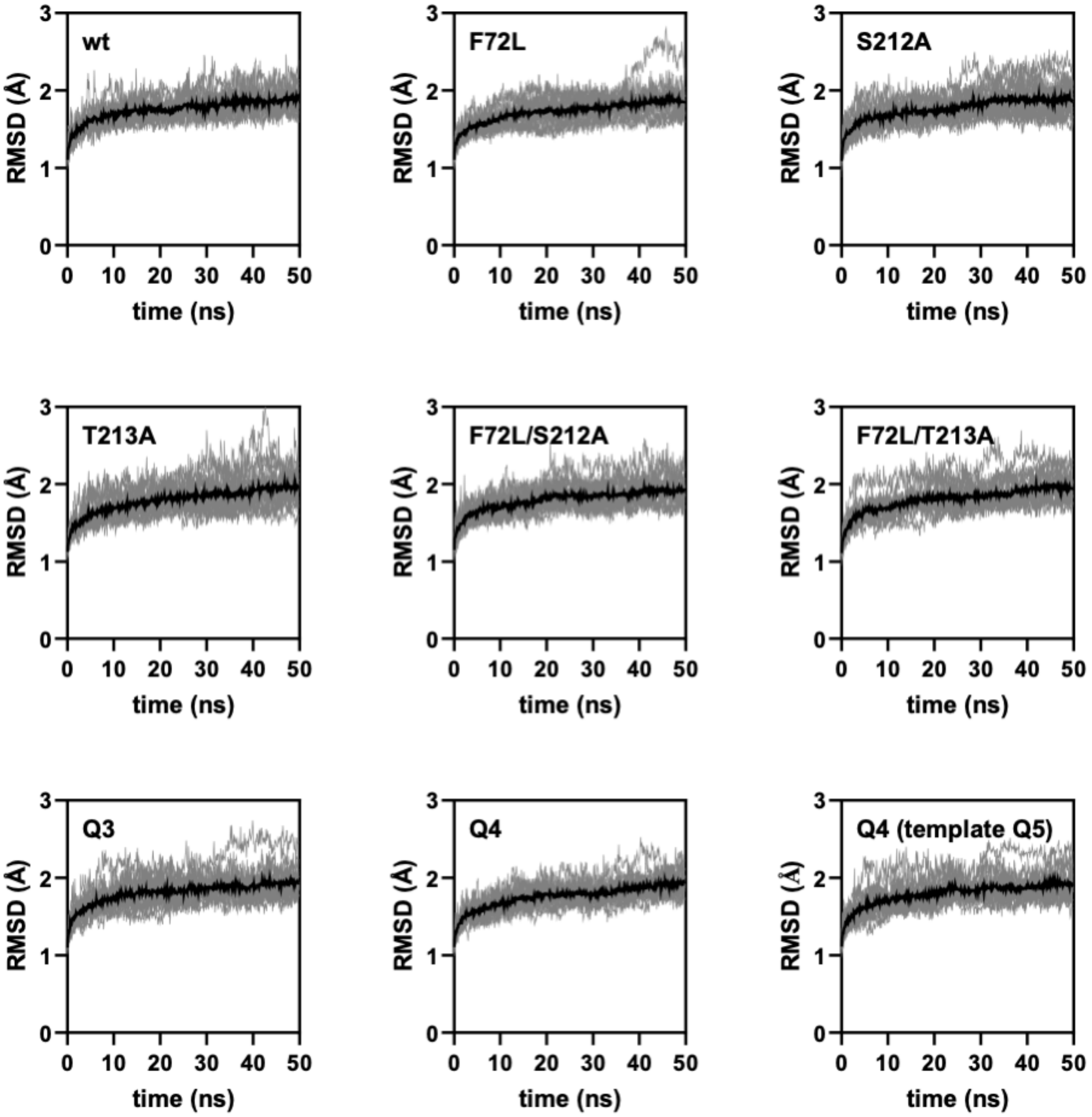


**Figure S7: MD simulations of the OXA-48 variants.**

RMSD plots for each variant indicate that the simulations are stable over 50 ns. The first 10 ns of each simulation were omitted from the subsequent analysis. The black line reflects the average of 20 independent replicates (grey). RMSD values were calculated based on $C_\alpha$ coordinates.

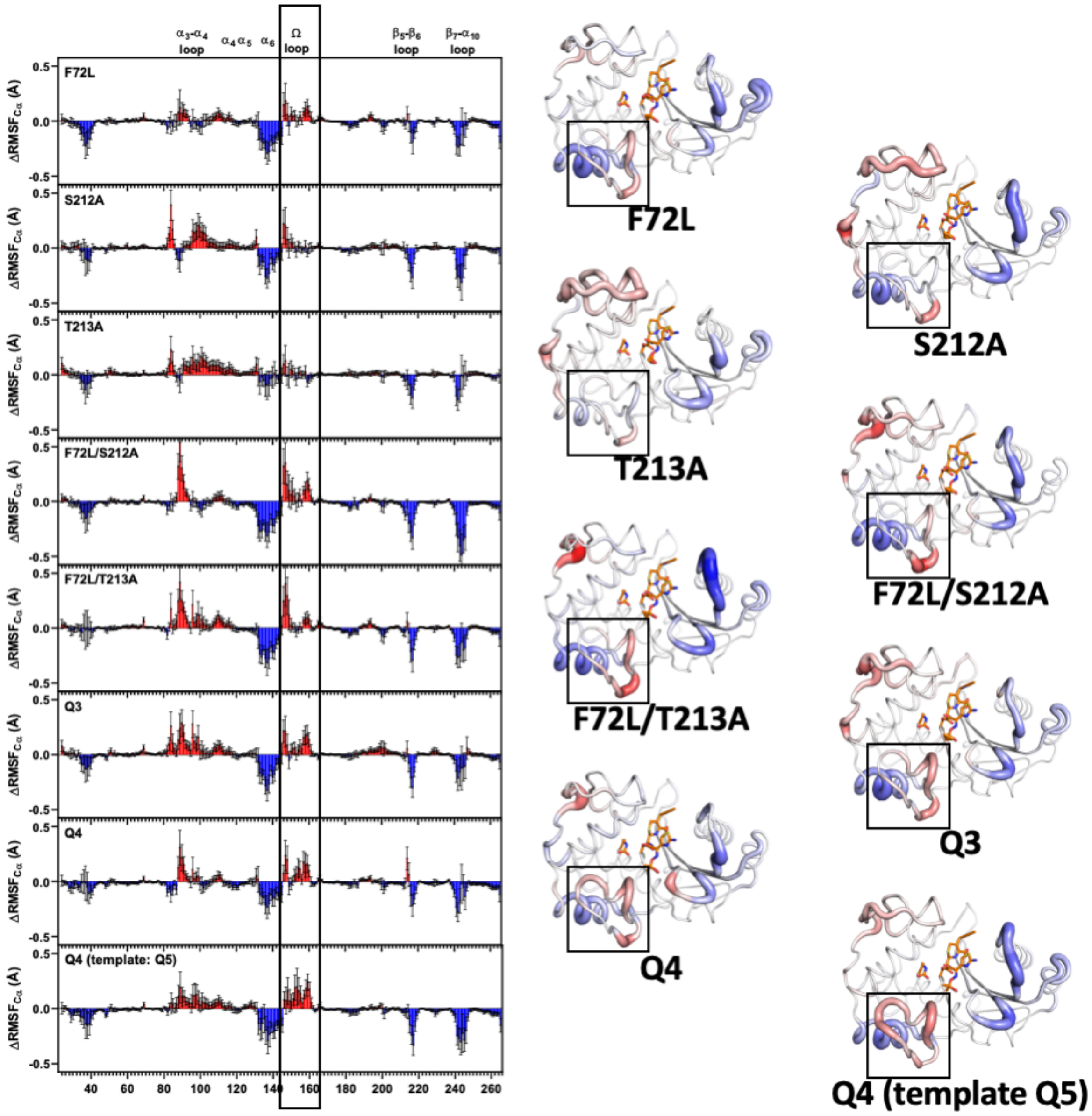


**Figure S8: Changes in per-residue $C_\alpha$ RMSF values compared to wtOXA-48.**
MD simulations reveal that F72L remodels flexibility in OXA-48, particularly in the Ω-loop. Residues that become more flexible compared to wtOXA-48 are shown in red, and those that become more rigid are shown in blue. In the structures, the tube thickness and color intensity are scaled to the overall change in dynamics. We note that the RMSF calculations tend to require more MD data to converge compared to the cluster analysis and dynamical correlations (Fig. S9 to S11), and are thus comparably more noisy (error bars reflect the standard error of the mean). Therefore, the dynamics of the system were subsequently analyzed using principal component (Fig. S9), cluster (Fig. S10), and dynamical correlation anaylsis (Fig. S11).

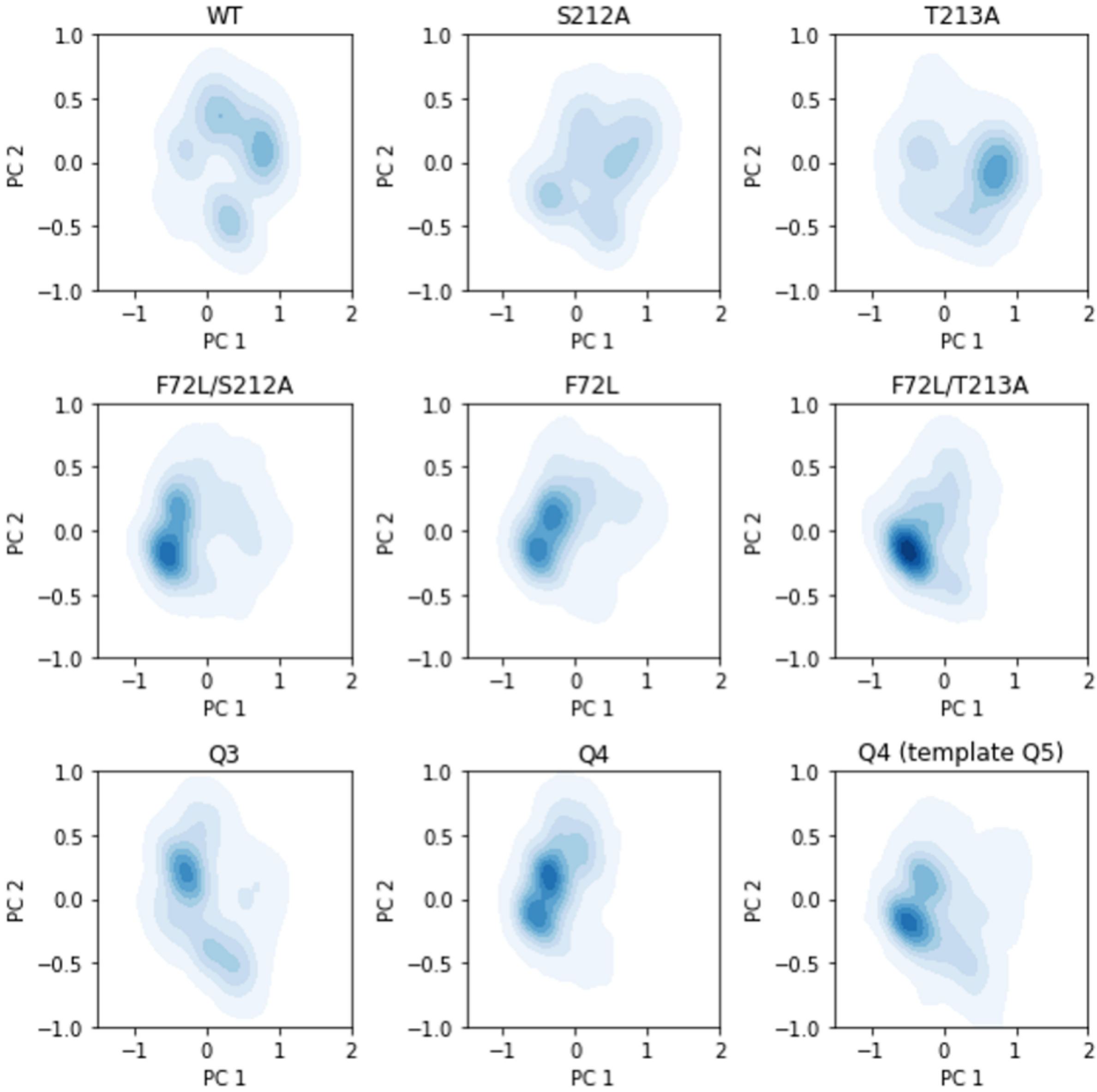


**Figure S9: Principal component (PC) analysis of the OXA-48 variants analysis of active-site loops (residues 96-106, 151-160, 213-218, 242-246).**

The F72L mutation changes the explored conformational landscape of OXA-48. The conformational landscape is only changed upon adding F72L, as revealed by PC analysis of the active-site loops (residues 96-106, 151-160, 213-218, 242-246). Notably, the S212A and T213A mutations only marginally affect the shape of the landscape in any genetic background.

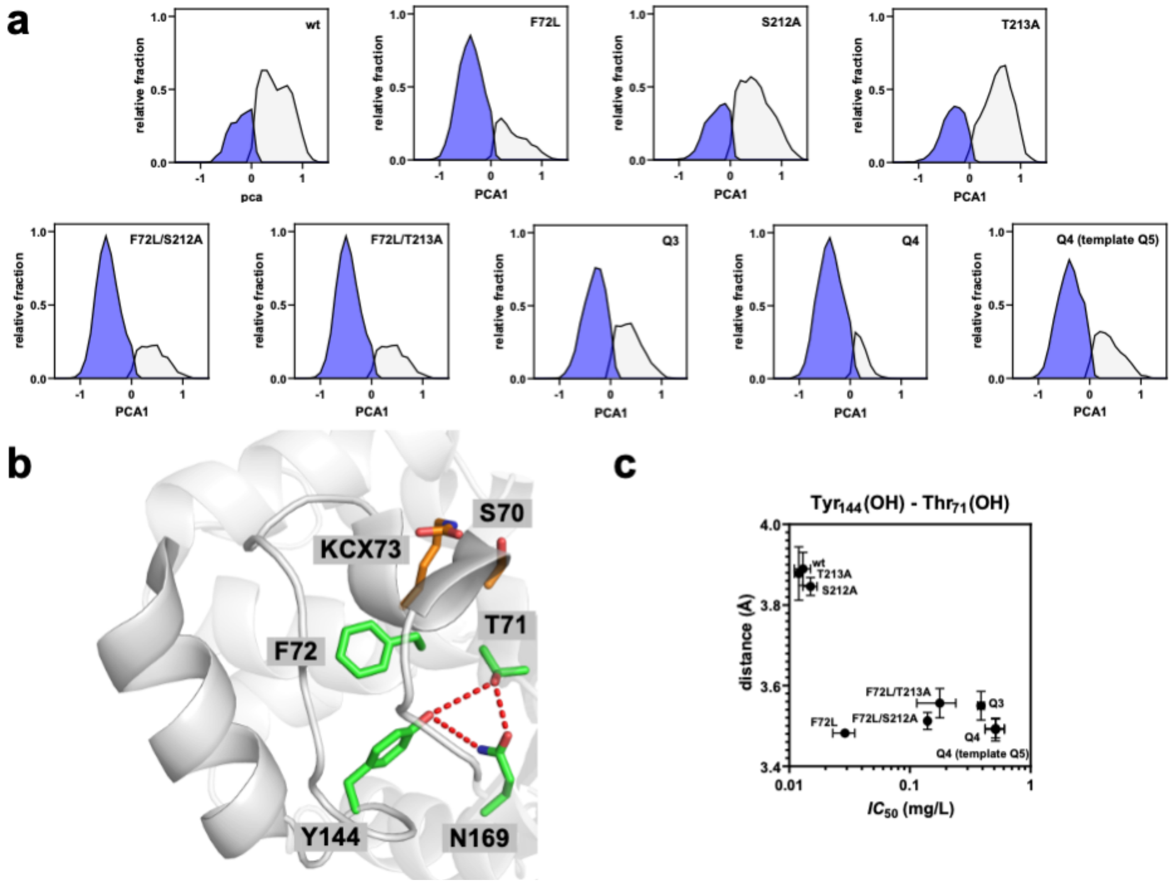


**Figure S10: Cluster analysis of the OXA-48 variants.**

**a.** Cluster analysis based on $C_\alpha$ RMSD of active-site loops (residues 96-106, 151-160, 213-218, 242-246) indicates an increase in population of a distinct state after the introduction of F72L. Trajectories (of all variants) were partitioned into two clusters using the k-means algorithm and projected onto PC1. We note that PC analysis (Fig. S9) indicates a broader conformational diversity than a two-state cluster model and that the two clusters thus represent a larger group of conformationally-similar sub-states. **b.** The distinct effect of F72L is exemplified by its effect on the H-bonding network (red dashed lines) comprising T71, Y144, and N169, **c.** which tightens up as soon as F72L is introduced, as indicated by the T71(OG)-Y144(OH) distance.

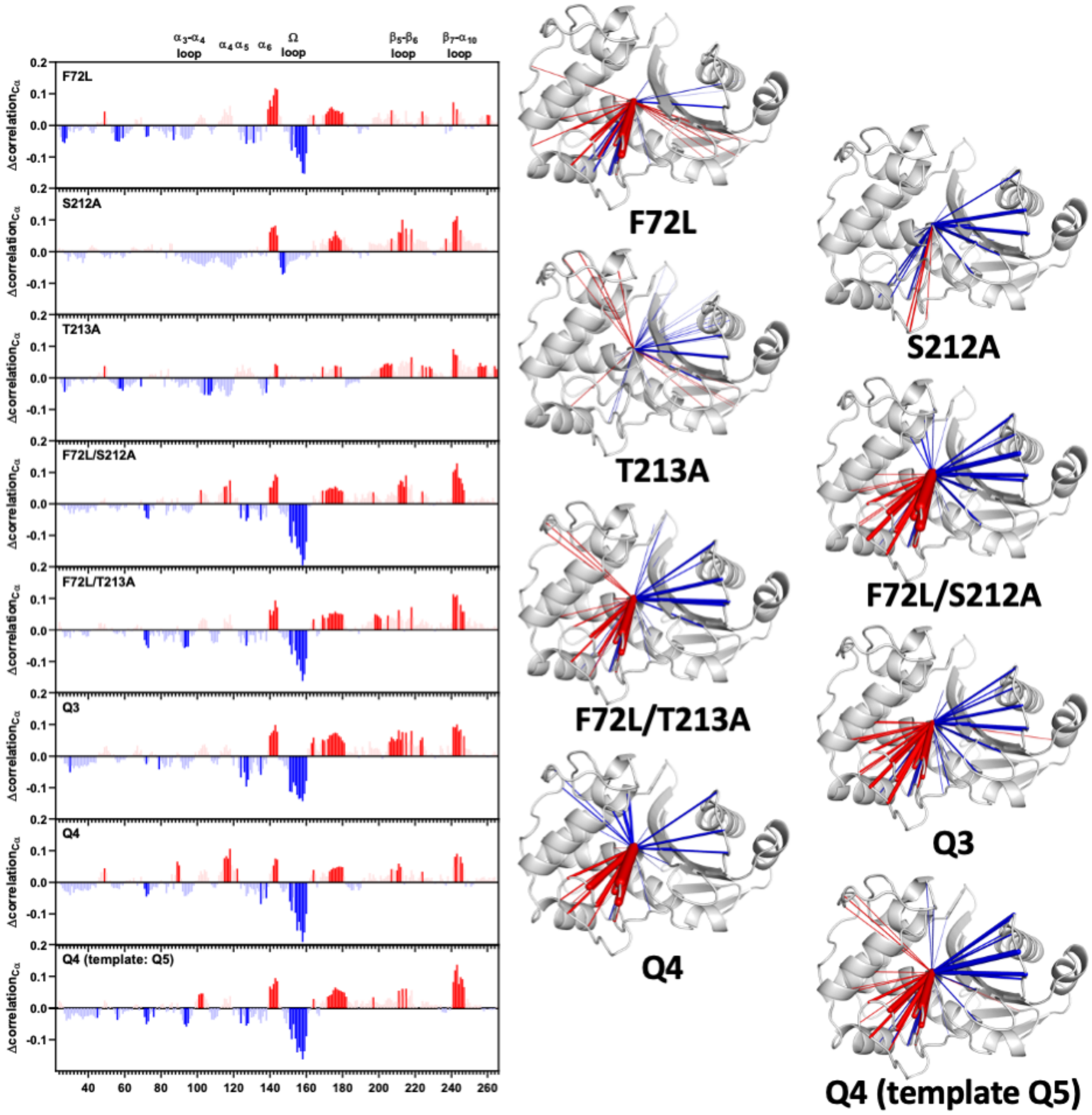


**Figure S11: Dynamical correlations in the OXA-48 variants.**

Changes in dynamical correlations of S70 with other protein residues are shown relative to wtOXA-48. The changes in correlations reveal distinct effects for F72L and the alanine mutations. F72L primarily reduced the correlation of S70 with the Ω-loop. In contrast, the alanine mutations increase correlation, particularly with the oxyanion-hole harboring half of the protein scaffold. Residues that become more correlated with S70 are highlighted in blue, and those that are less correlated in red. Only statistically significant changes compared to wtOXA-48 (T-test, α = 0.05, dark colors) are shown on the structures. The width of the lines in the structures corresponds to the magnitude of change in correlation.

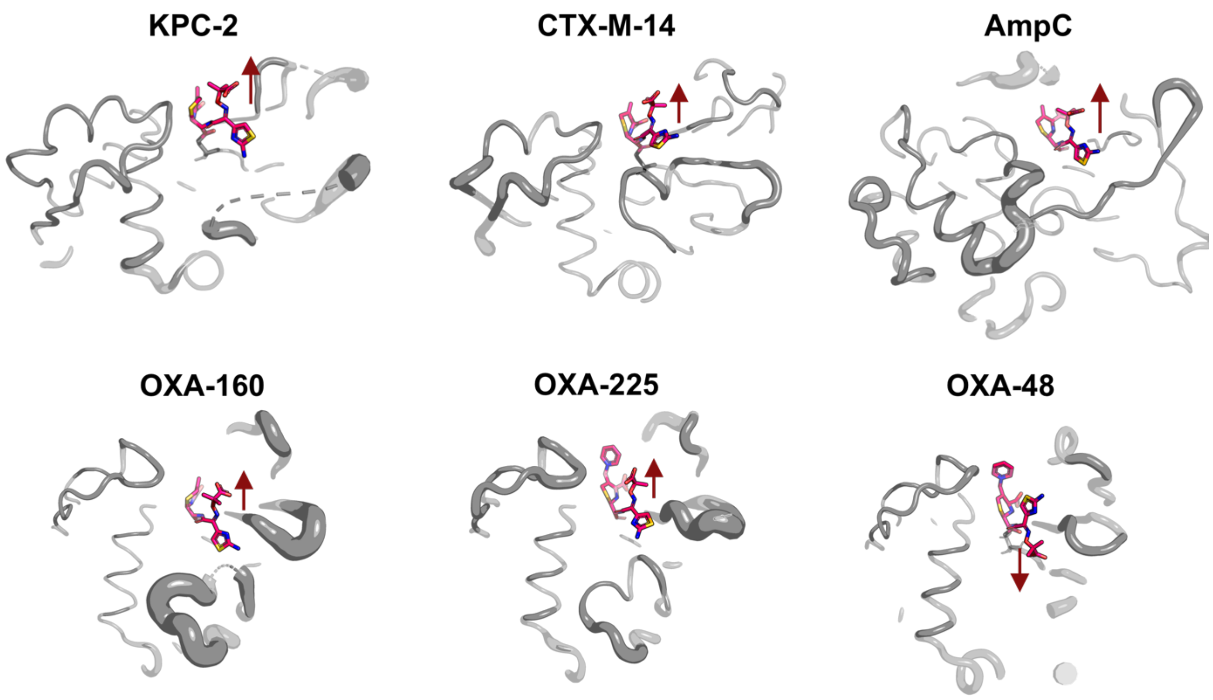


**Figure S12: Ceftazidime orientation in different β-lactamases.**

OXA-48 shows a distinct orientation of ceftazidime in the active site compared to other β-lactamases. Position of ceftazidime (pink) in the active site of KPC-2:E166Q (PDB ID: 6Z24[3]), CTX-M-14:E166A (PDB ID: 5U53[4]), AmpC (PDB ID: 1IEL[5]) as well as OXA-160:V130D (PDB ID: 4X56[6]), OXA-225:K82D (PDB ID: 4X55[6]) and OXA-48:P68A (PDB ID: 6Q5F[7]). The direction of the oxyimino group of ceftazidime is indicated by a red arrow, which points outwards from the active site for all depicted enzymes except for OXA-48:P68A.
